# *Ralstonia solanacearum* effector RipAF1 ADP-ribosylates host FBN1 to induce resistance against bacterial wilt

**DOI:** 10.1101/2022.04.04.487053

**Authors:** Wei Wu, Xuming Luo, Xinyu Chen, Leyi Wang, Kang Wang, Siyao Tian, Ziqunfang Tong, Tongxin Zhao, Xiaojing Fan, Tao Zhuo, Xun Hu, Huasong Zou

## Abstract

*Ralstonia solanacearum* (*Rso*) causes destructive bacterial wilt across a broad range of host plants by inducing jasmonic acid (JA) signaling while suppressing salicylic acid (SA) signaling pathways during disease development. In the present study, we show that *Rso* type III effector RipAF1 exerts a negative effect on bacterial virulence by subverting disease signaling in association with bacterial wilt. The ADP-ribosylation activity of RipAF1 was verified both *in vivo* and *in vitro*. Host fibrillin FBN1 was identified as a RipAF1-interacting protein that acted as a susceptible factor for bacterial wilt. In particular, RipAF1 directly ADP-ribosylates FBN1 at the E175/K207 residues, thus interfering with the mediation of disease signaling by FBN1. Together, these results suggest that RipAF1 exerts a role in defense induction by ADP-ribosylation of the susceptible factor FBN1 in the host plant.

## Introduction

The beta proteobacterium *Ralstonia solanacearum* (*Rso*) infects a wide range of host plant species, causing devastating bacterial wilt disease worldwide (Prior et al., 2016). *Rso* attaches to and penetrates into host roots by means of numerous infection mechanisms, including secretion of cell wall-degrading enzymes, biofilm formation, and production of exopolysaccharides (Genin and Denny, 2012). Furthermore, *Rso* injects a set of type Ⅲ effectors into host cells to modulate defense or susceptible reactions via a type Ⅲ secretion system (T3SS) (Coll and Valls, 2013). Through pan-genomic data analysis, 102 *Ralstonia* injected proteins (Rips) and 16 hypothetical T3E genes have been successfully identified from among 140 *Rso* species (Sabbagh et al., 2019) These *Rso* species possess an average of 64 T3Es, substantially more than *Xanthomonas campestris* and *Pseudomonas syringae* with averages of 23 and 31 T3Es, respectively (Dillon et al., 2019; Roux et al., 2015). As Rips exert diverse roles in pathogenicity, the pan-effectome of *Rso* strains results in a particular host width and specificity during infection (Landry et al., 2020).

The plant hormone jasmonic acid (JA) is a universal resistance signal that responds to necrotrophic pathogens in plants, whereas salicylic acid (SA) induces resistance to biotrophic and hemibiotrophic pathogens (Gomi, 2020). In addition, JA is closely involved in growth and developmental processes, such as stomatal opening, rubisco biosynthesis, and the absorption of nitrogen sources (Bertini et al., 2019; Jang et al., 2020; Taniguchi et al., 2014; Yoshitomi et al., 2016). *Rso* is a hemibiotrophic pathogen that induces antagonistic cross-talk between JA and SA signaling during infection of *Nicotiana benthamiana* (Nakano and Mukaihara, 2018, 2019a). After the initial *Rso* infection, JA production is activated, coupled with an inhibition of the SA-related signaling pathway (Nakano and Mukaihara, 2018). This progression suggests a complex signaling pathway is required for bacterial wilt development, within which JA is a susceptible signaling pathway and SA acts as a defense signaling pathway. JA and SA signaling are elaborately regulated, because during *Rso* infection, both are affected by a number of Rips, such as RipX, RipAL, RipI and RipAB (Nakano and Mukaihara, 2018; Qi et al., 2022; Sun et al., 2020; Zhuo et al., 2020).

One specific feature of *Rso* Rips is the abundance of genes that encode host catalyzing enzymes. This is well exemplified by six putative acetyltransferases, including RipJ, RipK, RipP1, RipP2, RipP3, and RipAE (Peeters et al., 2013; Sabbagh et al., 2019). RipAF1 is a putative ADP-ribosyltransferase (ADP-RT) that was identified as a Rip from the *Rso* RS1000 strain (Occhialini et al., 2005). Its homolog in other pathogenic bacteria, *avrpPhF*, was initially cloned from a 154-kb plasmid harbored by *Pseudomonas syringae* pv. *phaseolicola* (Tsiamis et al., 2000). Notably, the *avrpPhF* genetic locus comprises two open reading frames (ORFs), both of which are required for avirulence of the bacteria in bean cultivars carrying the R1 gene (Tsiamis et al., 2000). The two ORFs are involved in distinct biological processes; AvrpPhF ORF1 acts as a type Ⅲ chaperone, while ORF2 shares a structural similarity with the catalytic domain of diphtheria toxin, which has ADP-RT activity (Singer et al., 2004; Takada et al., 1995). AvrpPhF ORF2-mediated defense signaling has been well documented in association with the homologous HopF2 in *Pseudomonas syringae* pv. *tomato* DC3000 (Singer et al., 2004). HopF2 is able to mono ADP-ribosylate *Arabidopsis thaliana* MAP kinase kinase 5 (MKK5), thus inhibiting kinase activity that is essential for the activation of the MEKK1/MEKKs-MKK4/5-MPK3/6 cascade (Wang et al., 2010). Furthermore, the arginine and aspartic acid residues in positions 71 and 175, respectively, in HopF2 are necessary for the interaction with MKK5 to block MAP kinase activation (Wang et al., 2010). HopF2 also targets *Arabidopsis* RIN4 protein to promote *Pseudomonas syringae* virulence in *Arabidopsis* (Wilton et al., 2010).

Fibrillin family proteins are widely found in photosynthetic organisms ranging from cyanobacteria to vascular plants (Deruère et al., 1994; Hernández-Pinzón et al., 1999; Kessler et al., 1999). They are encoded by nuclear genetic loci and are closely involved in the biogenesis of chloroplast plastoglobules, stroma, thylakoids, and chromoplast fibrils (Kessler et al., 1999; Nacir and Bréhélin, 2013; Singh et al., 2012). Fibrillin mainly functions in plant growth, development, abiotic stress tolerance, and hormone signaling regulation (Singh and McNellis, 2011; Youssef et al., 2010). Knock down of the fibrillin gene *FBN1* in *Lycopersicon esculentum* resulted in increased susceptibility to the necrotrophic fungus *Botrytis cinerea* (Leitner-Dagan et al., 2006). Relative to their wild-type counterparts*, A. thaliana fib4* mutants and apple *fib4* RNA-interference transformants exhibited increased susceptibility to the bacterial pathogens *Pseudomonas syringae* pv. *tomato* and *Erwinia amylovora*, respectively (Singh et al., 2010).

RipAF1 is a conserved HopF2 family T3E protein in *Rso*, and it is localized to the cytoplasm, cell membrane, and nucleus of host cells (Jeon et al., 2020; Occhialini et al., 2005). The purpose of this work was to explore the role of T3E RipAF1 in *Rso*– host interactions. In addition to describing its negative role in virulence, we report that RipAF1 modifies the host susceptible factor FBN1 to subvert disease signaling pathways. These results not only provide a deeper and fuller understanding of RipAF1-mediated signaling pathways, but also highlight the role of ADP-ribosylation in *Rso*–host interactions.

## Results

### RipAF1 interferes with the expression of host JA and SA signaling marker genes

Based on the total repertoire analysis of T3E sequences (http://www.ralsto-t3e.org/), RipAF1 was determined to be present in representative strains from *Ralstonia* phylotypes I, IIA, and III, but absent in three phylotype IV strains, i.e., PSI07, BDBR229, and *R. syzygii*. Among the four representative phylotype IIB strains, RipAF1 is found in Po82 but absent from IPO1609, Molk2, and UW551 (Supplemental Figure S1, A). The existence of RipAF1 in the FJ1003 strain, isolated from a tobacco plant, was verified by PCR amplification. The full-length gene encodes a 350-amino-acid protein with high similarity to its homolog in GMI1000 encoded by Rsp0822, which is located on a plasmid. Based on phylogenetic analysis of inferred amino acid sequences, the evolution of RipAF1 is closely related to phylotype in that RipAF1 sequences of the same phylotype were confirmed to be monophyletic (Supplemental Figure S1, B). The RipAF1 sequence from FJ1003 was most closely related to GMI1000 RipAF1, which were grouped together with the RipAF1 sequences from phylotype I strains (Supplemental Figure S1, B). The RipAF1 sequence from FJ1003 has no transmembrane domain but possesses a putative mono-ADP-RT (mADP-RT) domain from positions 155 to 338 (Supplemental Figure S1, C). This domain shares 99% identity with the RipAF1 sequence from GMI1000 and 37% similarity with AvrPphF family HopF1 and HopF2 sequences from *Pseudomonas* (Supplemental Figure S1, D).

To examine the host response to RipAF1, the cloned RipAF1 from FJ1003 was transiently expressed in *N*. *benthamiana* driven by a 35S promoter in the binary vector pJL12. Compared with the empty vector control, the expression of RipAF1 induced weak visual chlorosis at 48 h post-infiltration, indicating that *N*. *benthamiana* exhibited a response to RipAF1 (Supplemental Figure S1, E). To determine whether RipAF1 affects the JA signaling pathway, the transcript levels of the JA marker genes *NbPDF1.2*, *NbOPR3,* and *NbLOX* were quantified. Compared to the empty vector control, transcript levels of *NbPDF1.2*, *NbOPR3,* and *NbLOX* were all significantly decreased in response to RipAF1 infection (Figure 1, A). *PDF1.2* was the gene with the most reduced expression, with an over^7^ 80% reduction relative to the control. For further confirmation of this suppression, the *PDF1.2* promoter was fused with a luciferase reporter and co-expressed with RipAF1-GFP. Compared with the GFP control, the expression of RipAF1-GFP remarkably inhibited luciferase activity, by over 62% (Figure 1, B and C). As salicylic acid (SA) signaling acts antagonistically to JA during bacterial wilt development, the transcript levels of SA signaling marker genes such as *NbICS1*, *NbPR1*, and *NbPR2* were studied. As expected, their expression levels were induced by RipAF1 (Figure 1, D).

**Figure 1.**
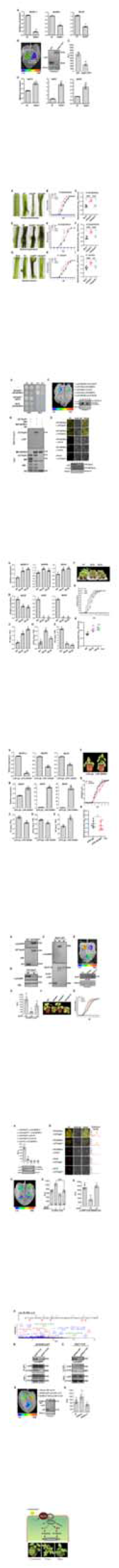
Effect of RipAF1 expression in *Nicotiana benthamiana* on jasmonic acid (JA) and salicylic acid (SA) signaling pathway marker genes. **A,** The reduced expression levels of the JA marker genes *NbPDF1.2*, *NbOPR3,* and *NbLOX* in *N*. *benthamiana* expressing RipAF1. Total RNA was isolated from leaves 48 h after agroinfiltration. Expression levels were determined by qRT-PCR analysis and normalized to that of the empty vector pJL12 (EV). Values are means ± SD (*n* = 3 biological replicates; Student’s *t*-test, \**p* < 0.05, \*\**p* < 0.01). **B,** Luciferase assays for the inhibition of *PDF1.2* promoter activity by RipAF1. *PDF1.2* promoter luciferase fusion (pro:PDF1.2-Luc) and RipAF1-GFP constructs were co-expressed in *N. benthamiana* leaves. Luciferase activity was measured with a CCD imaging system. The co-expression of pro:PDF1.2-Luc and GFP served as the negative control. The gels at right show the expression of GFP and RipAF1-GFP in western blots using α-GFP antibody. **C,** Quantitative assay of the luciferase signal of pro:PDF1.2-Luc. The signal was quantified with a microplate luminescence reader. Values are means ± SD (*n* = 6 biological replicates; Student’s *t*-test, \*\**p* < 0.01). **D,** The increased transcript levels of the SA signaling marker genes *NbICS1, NbPR1*, and *NbPR2* in *N*. *benthamiana* expressing RipAF1, performed as in B. Values are means ± SD (*n* = 3 biological replicates; Student’s *t*-test, \**p* < 0.05, \*\**p* < 0.01). All experiments were replicated three times with similar results, and representative results are shown.

### RipAF1 negatively contributes to bacterial virulence

To examine the contribution of RipAF1 to *R. solanacearum* virulence, *N. benthamiana* plants were inoculated with wild-type FJ1003, the *ripAF1* deletion mutant Δ*ripAF1*, and the complemented strain CΔ*ripAF1* via stem wound inoculation. The mutant *ΔripAF1* caused more severe wilt symptoms relative to wild-type FJ1003. At 7 days post-inoculation, the stem near the inoculation site of plants infected with *ΔripAF1* was completely dark and shrunken (Figure 2, A). A disease index was used to monitor bacterial wilt development, and mutant *ΔripAF1* induced a higher disease index until 10 days post-inoculation (Figure 2, B). Accordingly, *ΔripAF1* grew faster than wild-type FJ1003 in *N. benthamiana* (Figure 2, C). This increased virulence was also found in *Solanum lycopersicum* and *Capsicum annuum* plants; the mutant *ΔripAF1* increased disease severity, the disease index, and replication speed in both *S. lycopersicum* and *C. annuum* (Figure 2, D–I). To further validate the role of RipAF1 in virulence, a *ripAF1* deletion mutant generated from the GMI1000 strain was inoculated into *S. lycopersicum* and *C. annuum*. Compared with wild-type GMI1000, the *ripAF1* mutant caused more severe symptoms, increased disease indexes, and accelerated growth speeds in both *S. lycopersicum* and *C. annuum* plants (Supplemental Figure S2).

**Figure 2.**
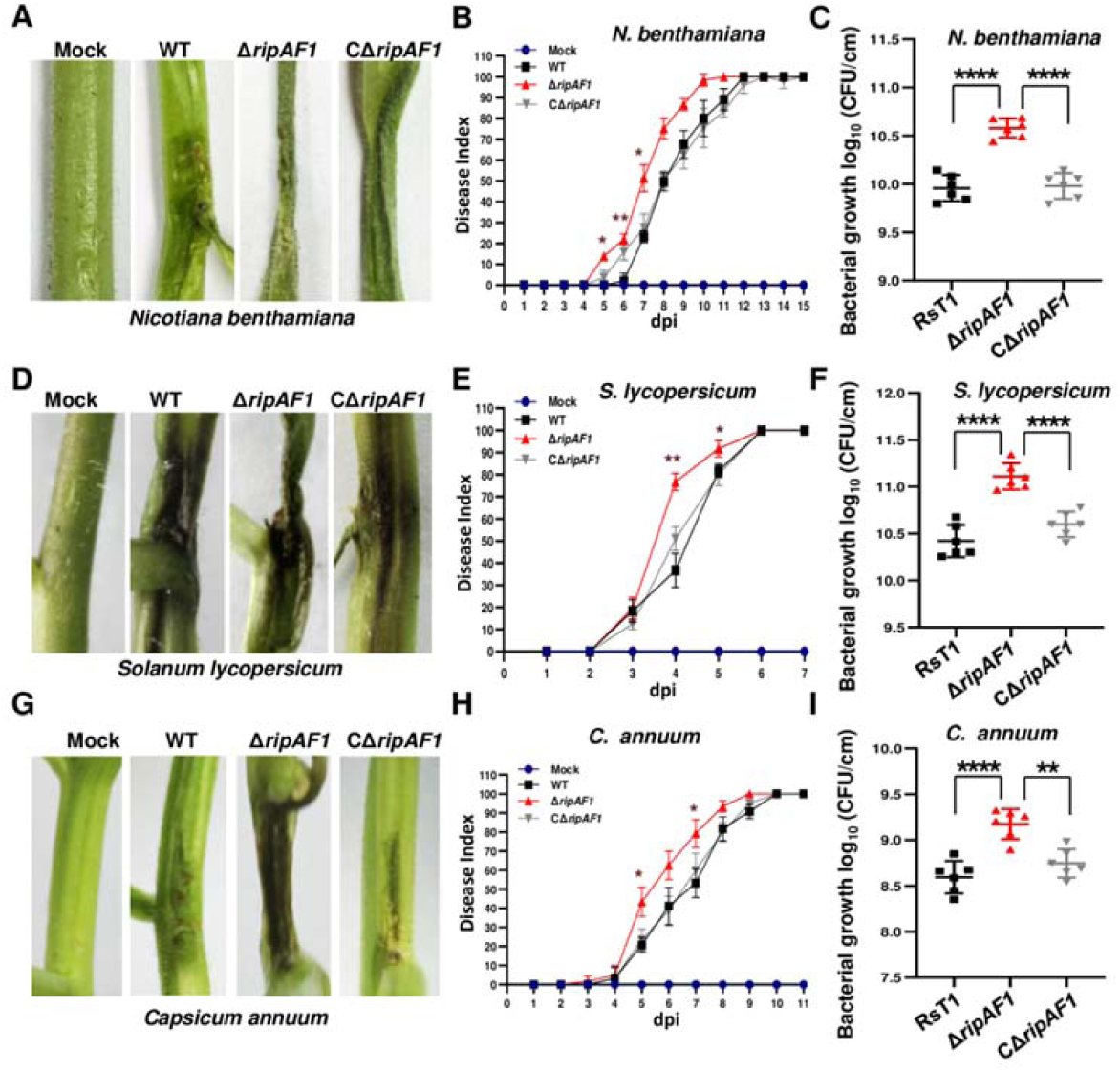
The enhanced virulence of the Δ*ripAF1* mutant of *Ralstonia solanacearum* FJ1003 on host plants. **A,** Disease symptoms of *Nicotiana benthamiana* plants inoculated with wild-type FJ1003, mutant Δ*ripAF1*, and the complemented strain CΔ*ripAF1.* Images were captured 7 d post-inoculation (dpi). The experiments were repeated three times with similar results, and representative results are shown. **B,** Progression of bacterial wilt on *N. benthamiana* plants inoculated with wild-type FJ1003, Δ*ripAF1*, and CΔ*ripAF1.* Disease severity was assessed after stem inoculation. Each time point represents the mean disease severity of 24 inoculated plants per treatment. Error bars represent the standard deviation of three independent experiments (two-way ANOVA, \**p* < 0.05, \*\**p* < 0.01). **C,** Growth of wild-type FJ1003, Δ*ripAF1*, and CΔ*ripAF1* in *N*. *benthamiana* plants. The stem of four-week-old *N*. *benthamiana* was injected with 100 μL of 10^6^ CFU/mL bacterial cell suspension. Plants were subjected to growth curve analysis at 3 dpi. Values are means ± SD (*n* = 6 biological replicates; Student’s *t*-test, \*\*\*\**p* < 0.0001). **D,** Disease symptoms in *Solanum lycopersicum* plants, characterized as in A. E, Progression of bacterial wilt on *S. lycopersicum* plants, characterized as in B. Disease severity was assessed after stem inoculation. **F,** Growth of the wild-type FJ1003, mutant Δ*ripAF1*, and CΔ*ripAF1* strains in *S. lycopersicum* plants, characterized as in C. Values are means ± SD (*n* = 6 biological replicates; Student’s *t*-test, \*\*\*\**p* < 0.0001). **G,** Disease symptoms in *Capsicum annuum* plants, characterized as in A. **H,** Progression of bacterial wilt on *C. annuum* plants, characterized as in B. Disease severity was assessed after stem inoculation. **I,** Growth of the wild-type FJ1003, Δ*ripAF1*, and CΔ*ripAF1* strains in *C. annuum* plants, characterized as in C. Values are means ± SD (*n* = 6 biological replicates; Student’s *t*-test, \*\**p* < 0.01, \*\*\*\**p* < 0.0001).

### RipAF1 interacts with host FBN1

To clone the host protein that interacted with RipAF1, yeast two-hybrid (Y2H) assays were conducted using RipAF1 as bait to screen the *N. benthamiana* cDNA library. A positive clone was identified, and it harbored a 1162-bp DNA fragment corresponding to the fibrillin gene *FBN1a*. Except for a 966-bp opening read frame, the clone contained a 61-bp 5′ terminal untranslated region (UTR) and a 135-bp 3′ terminal UTR. The Y2H experiment was repeated, followed by cloning the 966-bp ORF of *NbFBN1a* into the pGADT7 and pGBKT7 vectors. The positive interaction was verified after yeast AH109 was co-transformed with either pGADT7-RipAF1/pGBKT7-NbFBN1a or pGADT7-NbFBN1a/pGBKT7-RipAF1 (Figure 3, A; Supplemental Figure S3, A). In a MBP pull down assay, MBP-NbFBN1a successfully pulled down GST-RipAF1. In the control lane, MBP tag did not pull down RipAF1, and the MBP-NbFBN1a construct was unable to pull down GST tag (Figure 3, B). *N. benthamiana* has two copies of *FBN1*, named *NbFBN1a* and *NbFBN1b* (*NbFBN1a/b* hereafter). Each of the inferred FBN1 proteins has 321 amino acids, sharing 95% identity with 16 amino acid differences (Supplemental Figure S3, B). The extremely high identity suggests that NbFBN1a and NbFBN1b are functionally redundant in *N. benthamiana*. Fluorescence was mainly observed in the plastid, and a slight signal from the plasma membrane was detected when NbFBN1a/b-GFP was expressed in *N. benthamiana* (Supplemental Figure S3, C). Transient expression assays using *N. benthamiana* protoplasts also indicated that NbFBN1a/b was mainly located in the plastid (Supplemental Figure S3, D). Expression of RipAF1-mcherry in *N. benthamiana* revealed that RipAF1-mcherry was localized to the cytoplasm, membrane, and nucleus (Supplemental Figure S3, E). Additionally, merged yellow fluorescence was observed in the plasma membrane when RipAF1-mcherry was transiently co-expressed with NbFBN1a/b-GFP (Supplemental Figure S3, E). This demonstrated that RipAF1 and NbFBN1a/b were co-localized to the cell membrane. The interaction between RipAF1 and FBN1a/b *in vivo* was first verified by split luciferase assays (Figure 3, C). Then, the interactions at the plasma membrane were confirmed by BiFC assays (Figure 3, D). Meanwhile, the membrane localization was indicated by using the membrane marker PIP2A-mcherry. Merged orange fluorescence was localized to the cell membrane when membrane-anchored PIP2A- mcherry was co-expressed with BiFC constructs (Supplemental Figure S3, F).

**Figure 3.**
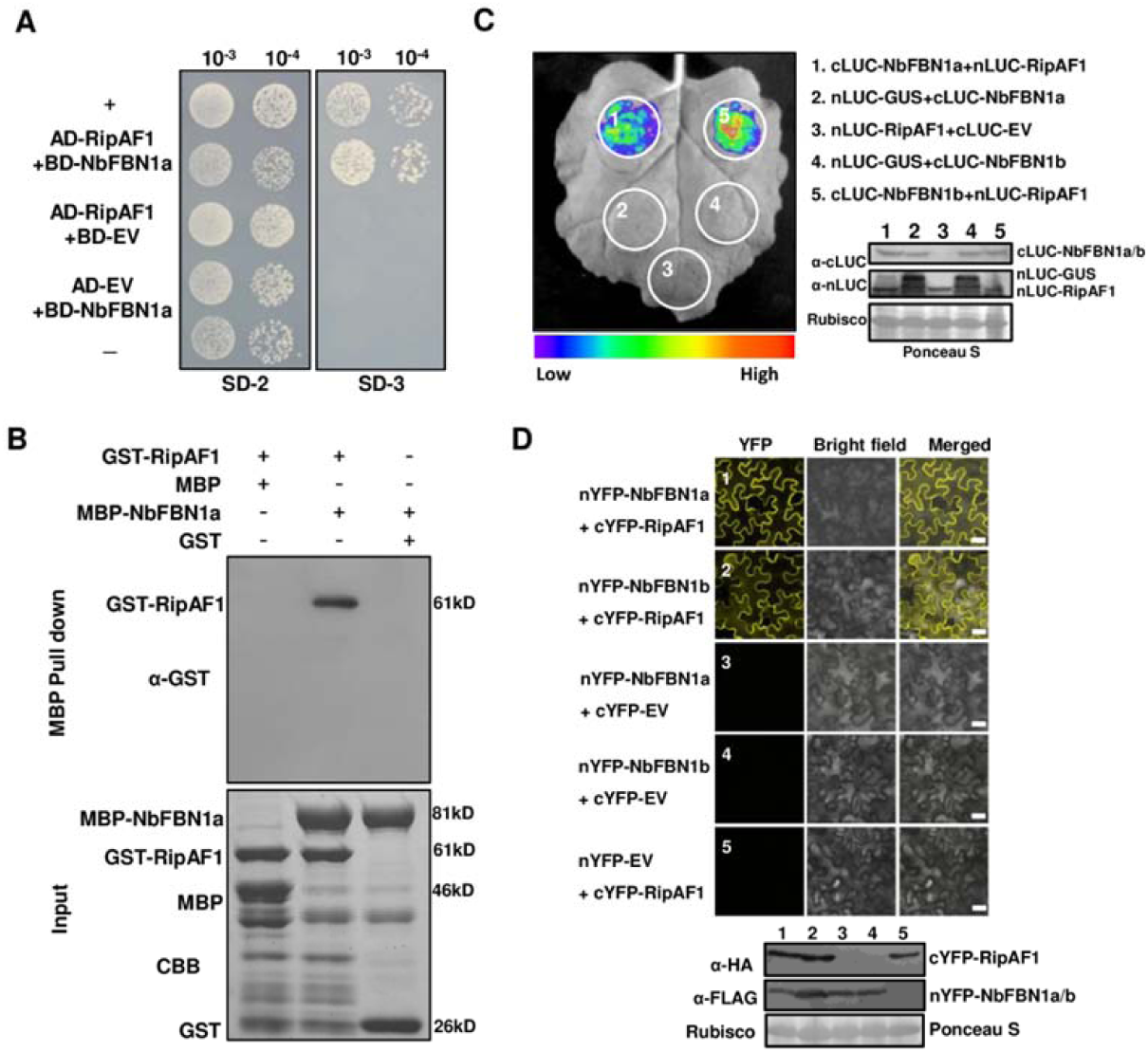
RipAF1 interactions with *Nicotiana benthamiana* FBN1 *in vitro* and *in vivo*. **A,** Yeast two-hybrid analysis showing RipAF1 interacts with NbFBN1a. AD-RipAF1 and BD-NbFBN1a were co-transformed into yeast cells and screened on synthetic dextrose media lacking leucine and tryptophan (SD/-Leu-Trp). A single yeast colony was then selected for serial dilution and subsequent culture on SD/-Leu-Trp (SD-2) and SD/-Leu-Trp-His (SD-3) to examine the potential interaction. Yeast co-transformed with pGADT7-T and pGBKT7-53 served as the positive control (+), while yeast co-transformed with pGADT7-T and pGBKT7-lam served as the negative control (-). EV, empty vector. **B,** MBP pull-down assay of the interaction between RipAF1 and NbFBN1a. The recombinant GST-RipAF1 and MBP-NbFBN1a were subjected to MBP pull-down analysis. MBP and GST proteins served as the negative control. The pulled down GST-RipAF1 was detected by anti-GST immunoblotting. The experiment was replicated three times with similar results. **C,** Split luciferase assay for the interaction of RipAF1 with NbFBN1a/b. *N. benthamiana* leaves were co-infiltrated with *Agrobacterium* carrying the 35S:RipAF1*-*nLUC and 35S:cLUC*-*NbFBN1a/b constructs. Images of chemiluminescence were obtained after application of 0.5 μM luciferin 48 h post-infiltration. Western blotting showed the expression of the respective proteins. Similar results were observed in three biological replicates. **D,** Split YFP assay for the interaction of RipAF1 and NbFBN1a/b, performed as in C. RipAF1 was fused with cYFP at the C terminus, and NbFBNs was fused with nYFP at the N terminus. The images were observed under a confocal microscope at 2 days post-agroinfiltration. Scale bar, 25 μm. Western blotting was used to confirm the expression of the respective proteins.

Notably, NbFBN1 homologs were found in *S. lycopersicum* and *C. annuum.* Their amino acid sequences shared over 61% identity with NbFBN1a/b. In the phylogenetic analysis of FBN1 from *N. benthamiana*, *S. lycopersicum*, *C. annuum*, and *A. thaliana*, FBN1 from *S. lycopersicum* and *C. annuum* showed a close orthologous relationship with NbFBN1a/b (Supplemental Figure S4, A). The interactions between RipAF1 and FBN1 from *S. lycopersicum* and *C. annuum* were verified by split luciferase assays (Supplemental Figure S4, B). This was consistent with RipAF1 being able to interact with FBN1 from various solanaceous plants, indicating that host FBN1 is the key target mediated by RipAF1 to affect virulence.

### Overexpression of NbFBN1a increases JA signaling and suppresses SA signaling

To study the role of *NbFBN1* in bacterial wilt development, NbFBN1a-GFP fusion constructs were transiently overexpressed in *N. benthamiana*. The expression of NbFBN1a-GFP did not cause any visible phenotype. However, compared with the GFP control, NbFBN1a-GFP significantly induced JA signaling pathway marker genes *NbPDF1.2*, *NbOPR3,* and *NbLOX*. The transcript level of *PDF1.2* was the most induced, increasing 5.5-fold (Supplemental Figure S5, A). In contrast, the expression levels of SA signaling pathway marker genes *NbICS1*, *NbPR1*, and *NbPR2* were all suppressed (Supplemental Figure S5, B). The induction of *NbPDF1.2* was additionally verified by co-expression of the *NbPDF1.2* promoter luciferase fusion construct with NbFBN1a-GFP. Compared with the GFP control, NbFBN1a-GFP significantly induced luciferase activity, verifying that transient overexpression of NbFBN1a induced the expression of *NbPDF1.2* (Supplemental Figure S5, C and D).

To confirm the altered signaling pathways, transgenic *N. benthamiana* plants expressing *NbFBN1a-GFP* were generated. Two independent homozygotes, lines 25 and 29, respectively, were chosen for further analysis after the overexpression of NbFBN1a was verified by qRT-PCR, western blot, and GFP fluorescence assays (Supplemental Figure S6). In the two transgenic lines, the transcript levels of *NbPDF1.2*, *NbOPR3*, and *NbLOX* were enhanced, while the transcript levels of SA signaling marker genes *NbICS1*, *NbPR1*, and *NbPR2* were reduced (Figure 4, A and B). In addition, the biosynthesis of JA, jasmonoyl-isoleucine (JA-Ile), and SA was quantified in the transgenic plants. In both lines 25 and 29, the biosynthesis of JA and JA-Ile was increased, while SA biosynthesis was reduced (Figure 4, C-E). Transgenic plants were examined for their resistance to bacterial wilt. The disease indexes of both transgenic lines were higher than that of wild-type plants, suggesting that lines 25 and 29 were more sensitive to *Rso* than wild-type *N. benthamiana* (Figure 4, F and G). Consistent with the increased susceptibility observed, *Rso* FJ1003 replicated faster in NbFBN1a-GFP transgenic plants (Figure 4, H).

**Figure 4.**
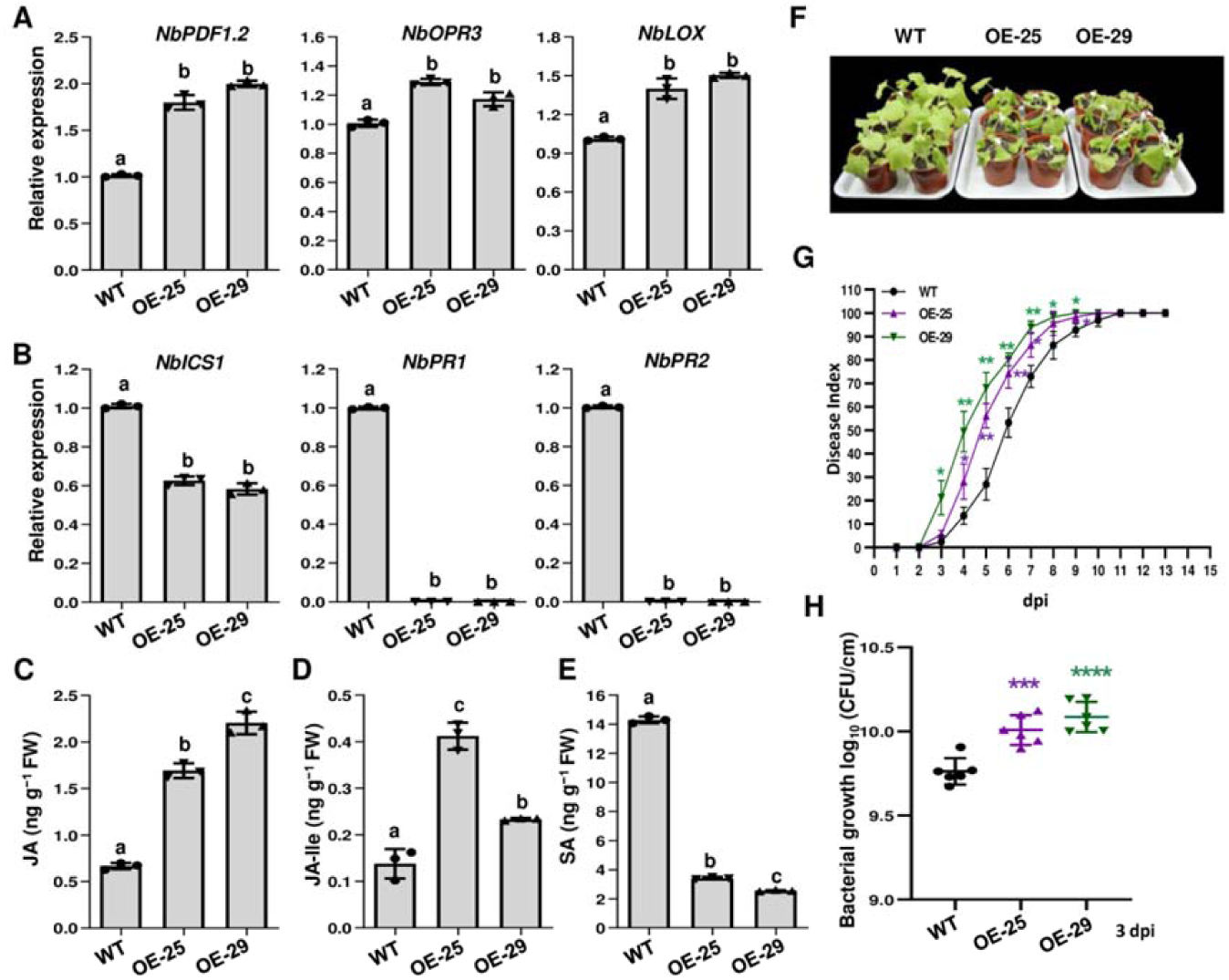
*NbFBN1a* transgenic *Nicotiana benthamiana* was more sensitive to bacterial wilt. **A,** The expression levels of jasmonic acid (JA) marker genes were enhanced in *NbFBN1a* transgenic plants. Total RNA was isolated from four-week-old transgenic *NbFBN1a*-overexpression lines OE-25 and OE-29. The transcript level for each gene in transgenic plants was compared with that in wild-type plants to monitor the expression change. Columns labeled with the same letter indicate means were not significantly different. Values are means ± SD (*n* = 3 biological replicates; ANOVA, *p* < 0.01). **B,** The expression levels of salicylic acid (SA) marker genes were reduced in *NbFBN1a* transgenic plants, assayed as in A. **C, D, and E** *NbFBN1a* transgenic plants showed increased contents of JA **(C)** and jasmonoyl-isoleucine (JA-Ile) **(D)** and reduced content of SA **(E)**. Phytohormone levels were quantified by HPLC-MS/MS. Values are means ± SD (*n* = 3 biological replicates). Columns labeled with different letter indicate significantly different means (ANOVA, *p* < 0.01). **F,** Wilt symptoms in wild-type and *NbFBN1a* transgenic plants inoculated with *Ralstonia solanacearum* FJ1003. Images were captured 7 d post-inoculation (dpi). All experiments were replicated three times with similar results, and representative results are shown. **G,** Disease index of *NbFBN1a* transgenic plants inoculated with *R*. *solanacearum* FJ1003. Each time point represents the mean disease severity of 24 inoculated plants per treatment. Error bars represent the standard deviation from three independent experiments (ANOVA, \**p* < 0.05, \*\**p* < 0.01). **H,** Growth of the FJ1003 strain in *NbFBN1a* transgenic plants. Plants stems were collected for growth curve analysis at 3 dpi with 100 μL of 10^6^ CFU/mL bacterial cell suspension. Values are means ± SD (*n* = 6 biological replicates; Student’s *t*-test, \*\*\**p* < 0.001, \*\*\*\**p* < 0.0001).

### Knock down of *NbFBN1* promotes *N*. *benthamiana* resistance to bacterial wilt

The knock down of *NbFBN1* in *N*. *benthamiana* was conducted by using tobacco rattle virus (TRV)-induced gene silencing. The pTRV:*gfp* construct carrying a 358-bp fragment of the *gfp* gene was used as a control. Compared with the *gfp-*silenced control, the transcript level of *NbFBN1* was decreased in pTRV:*NbFBN1* plants, indicating *NbFBN1a/b* was successfully knocked down (Supplemental Figure S7, A). The knock down of *NbFBN1* suppressed plant growth, especially root development (Supplemental Figure S7, B). In the *NbFBN1*-silenced plants, the transcript levels of JA marker genes *NbPDF1.2*, *NbOPR3*, and *NbLOX* were reduced (Figure 5, A). The decreased expression of *NbPDF1.2* was further verified by expression of *NbPDF1.2* promoter luciferase fusion protein in *NbFBN1*-silenced plants (Supplemental Figure S7, C). The expression levels of three SA signaling marker genes, *NbICS1*, *NbPR1*, and *NbPR2*, were each enhanced (Figure 5, B). Accordingly, JA and JA-Ile levels were significantly decreased, and SA was increased in *NbFBN1*-silenced plants (Figure 5, C–E). The inoculation of the FJ1003 strain into *NbFBN1*-silenced plants induced retarded disease development, and the disease index was lower than that of pTRV:*gfp* control plants (Figure 5, F and G). The replication of FJ1003 was decreased in *NbFBN1-*silenced plants (Figure 5, H). The increased resistance to bacterial wilt of knock-down plants suggested that *NbFBN1* serves as a bacterial wilt susceptibility gene.

**Figure 5.**
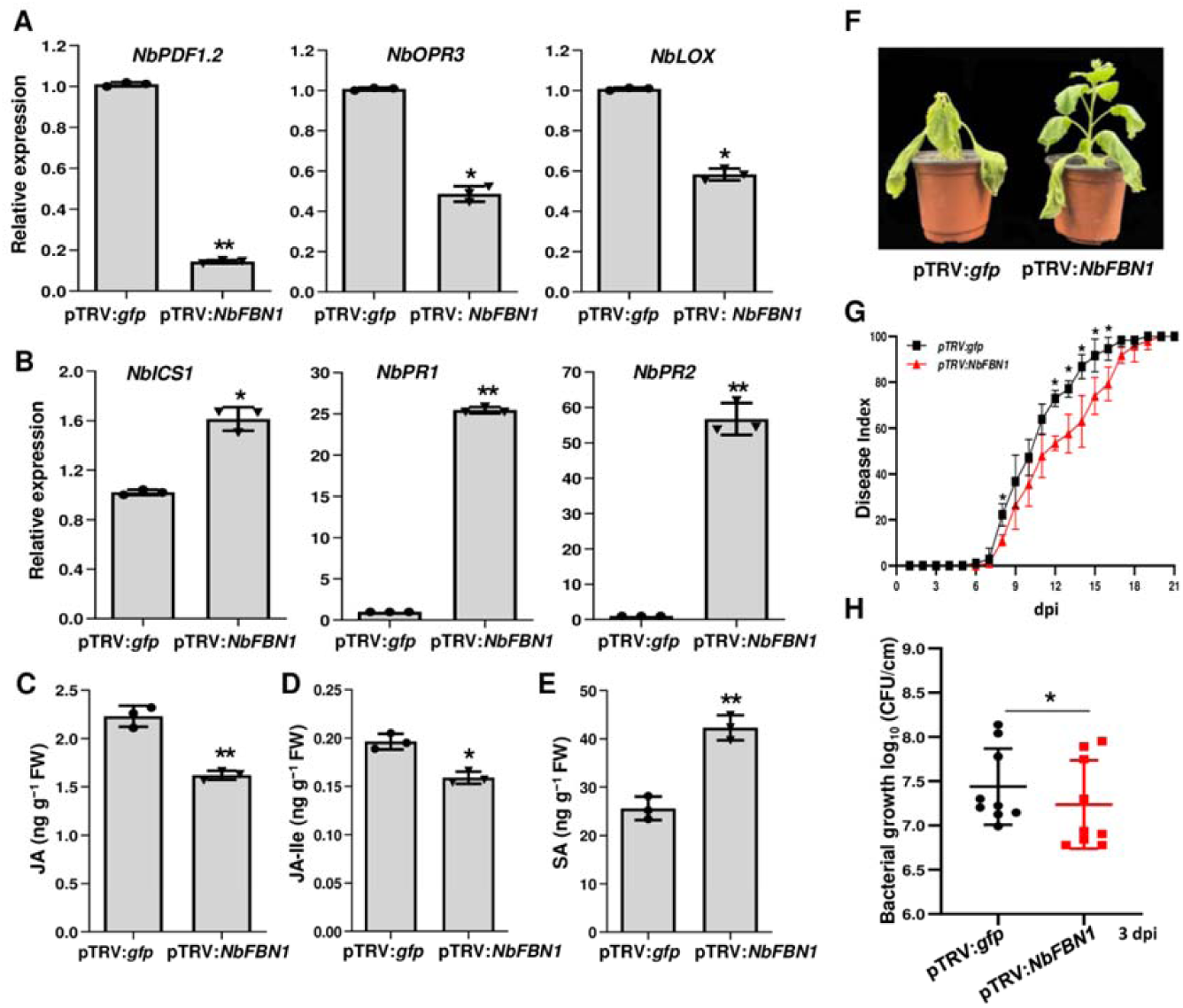
Knock down of *NbFBN1* in *Nicotiana benthamiana* resulted in enhanced resistance to bacterial wilt. **A,** Expression levels of jasmonic acid (JA) signaling marker genes were reduced in pTRV:*NbFBN1* plants. Total RNA was isolated from the new upper leaves when photobleaching was observed in phytoene desaturase (*PDS*)-silenced plants (efficiency control). The transcript level in plants transformed with the tobacco rattle virus pTRV:*gfp* construct was used as a control to monitor expression change. Values are means ± SD (*n* = 3 biological replicates; Student’s *t*-test, \**p* < 0.05, \*\**p* < 0.01). **B,** The expression levels of salicylic acid (SA) signaling marker genes were enhanced in pTRV:*NbFBN1* plants. The analyses were performed as in **A**. **C, D, and E** The reduced biosynthesis of JA **(C)** and jasmonoyl-isoleucine (JA-Ile) **(D)** and increased biosynthesis of SA **(E)** in pTRV:*NbFBN1* plants. Phytohormones were quantified by HPLC-MS/MS. Values are means ± SD (*n* = 3 biological replicates; Student’s *t*-test, \**p* < 0.05, \*\**p* < 0.01). **F,** The attenuated wilt symptom of pTRV:*NbFBN1* plants. Images of representative plants were captured 9 d post-inoculation (dpi). The experiments were replicated three times with similar results, and representative results are shown. **G,** Disease index of pTRV:*NbFBN1* and pTRV:*gfp* plants inoculated with the *Ralstonia solanacearum* FJ1003 strain. Each time point represents the mean disease severity of 24 inoculated plants per treatment. Error bars represent the standard deviation from three independent experiments (ANOVA, \**p* < 0.05). H, Growth of *R*. *solanacearum* FJ1003 in pTRV:*NbFBN1* and pTRV:*gfp* plants. The stems of plants were injected with 100 μL of 10^6^ CFU/mL bacterial cell suspension. Plants were subjected to growth curve analysis at 3 dpi. Error bars represent means ± standard deviation (Student’s *t*-test, \**p* < 0.05, *n* = 9).

### The Arg191/Asp310 residues in RipAF1 are necessary for mono-ADP-ribosylation activity and contribute to virulence

As RipAF1 possesses a mADP-RT domain, an anti–pan-ADP-ribose binding reagent (anti-panADPR) was used to examine whether RipAF1 has mADP-RT activity. GST-tagged RipAF1 was first expressed in *Escherichia coli* strain BL21 and purified for subsequent analysis. The mADP-RT activity of RipAF1 was successfully detected by using anti-panADPR. A specific activity band was observed in GST-RipAF1 samples, but not in the GST tag control (Figure 6, A). The mADP-RT domain of RipAF1 showed 37% identity with that of HopF2, whose Arg71 and Asp175 residues are critical for its mADP-RT activity (Supplemental Figure S8, A–D). The corresponding residues in RipAF1 are Arg191 and Asp310, which are located in the exposed surface of the predicted 3D molecular structure (Supplemental Figure S8, E). To examine the roles of these two residues in mADP-RT activity, Arg191 and Asp310 were both mutated to alanine, thus generating RipAF1^R191A/D310A^ (RipAF1^2A^). RipAF1^2A^ showed the same subcellular localization as wild-type RipAF1. Fluorescence was observed in the cell membrane, cytoplasm, and nucleus when RipAF1^2A^-GFP was expressed in *N. benthamiana* (Supplemental Figure S8, F). Most importantly, the mADP-RT activity of RipAF1^2A^ was impaired when it was expressed in *E. coli* (Figure 6, B). Immunoprecipitation of RipAF1^2A^-GFP from in *N. benthamiana* revealed that its mADP-RT activity was completely lost *in planta* (Figure 6, C). Compared with wild-type RipAF1, RipAF1^2A^ remarkably reduced the suppression effect on *PDF1.2* promoter activity (Figure 6, D and E). Furthermore, expression of RipAF1^2A^ in *ΔripAF1* did not restore the virulence phenotype to that of wild-type FJ1003 (Figure 6, F and G). Therefore, the Arg191 and Asp310 residues of RipAF1 were essential for mADP-RT activity and contributed to its virulence.

**Figure 6.**
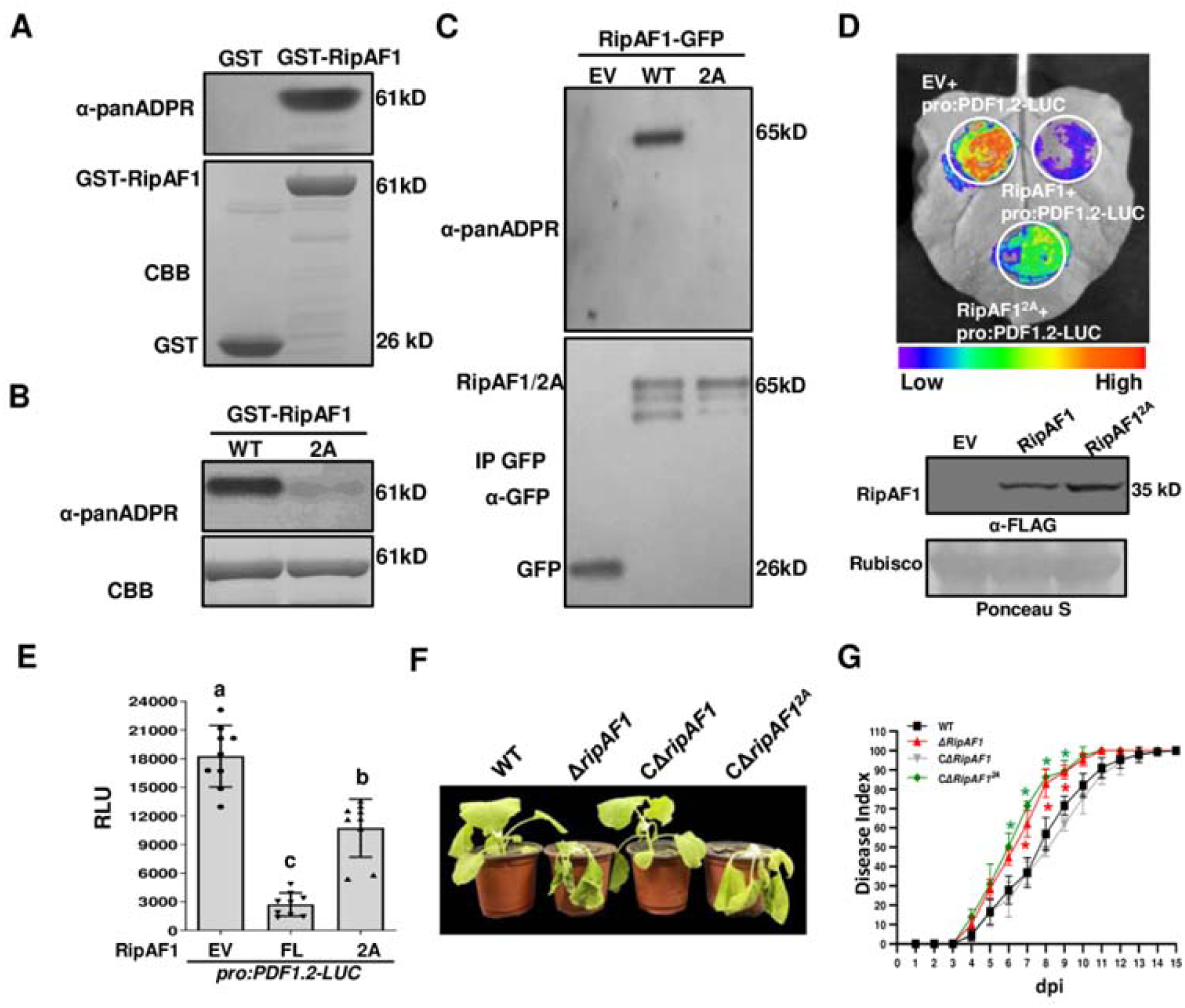
The Arg191/Asp310 residues in RipAF1 were critical for mono-ADP-ribosyltransferase activity. **A,** RipAF1 displayed mono ADP-ribosyltransferase (mADP-RT) activity *in vitro*. The mADP-RT activity of recombinant GST-RipAF1 was examined using anti-panADPR reagent. GST served as the negative control. Protein abundance was revealed by Coomassie brilliant blue (CBB) staining. Similar results were obtained from three independent replicates. **B,** Mutation of R191 and D310 in RipAF1 to alanine impaired mADP-RT activity. GST-RipAF1 and GST-RipAF1^2A^ (R191 arginine and D310 aspartic were substituted with alanine) were subjected to mADP-RT activity analysis. Protein abundance was indicated by CBB staining. Similar results were obtained from three independent replicates. **C,** The complete loss of mADP-RT activity of RipAF1^2A^ *in vivo*. RipAF1-GFP and RipAF1^2A^-GFP were expressed in *Nicotiana benthamiana,* immunoprecipitated with anti-GFP agarose beads and subjected to mADP-RT activity analysis. GFP was used as the negative control. The experiment was replicated three times with similar results. **D,** RipAF1^2A^ showed an attenuated suppression effect on *PDF1.2* promoter activity relative to wild-type RipAF1. The pro:PDF1.2-Luc construct was co-expressed with RipAF1-FLAG or RipAF1^2A^-FLAG in *N. benthamiana* leaves. Luciferase activity was measured with a CCD imaging system. The co-expression of pro:PDF1.2-Luc with empty vector was used as the negative control. The gels below show the expression of respective proteins. **E,** Quantitative assays of *PDF1.2* promoter activity when co-expressed with RipAF1 or RipAF1^2A^. Quantification of the luciferase signal was performed with a microplate luminescence reader. Values are means ± SD (*n* = 9 biological replicates). Columns labeled with different letters represent significantly different means (ANOVA with Tukey’s test, *p* < 0.01). **F,** RipAF1^2A^ was unable to rescue the virulence alteration of mutant Δ*ripAF1.* Disease symptoms of plants inoculated with the following *Ralstonia solanacearum* strains were examined: wild-type FJ1003, mutant Δ*ripAF1*, and the complemented strain CΔ*ripAF1* or CΔ*ripAF1^2A^*. Images were captured 7 d post-inoculation. The experiments were replicated three times with similar results, and representative results are shown. **G,** Progression of bacterial wilt on *N. benthamiana* plants inoculated with wild-type FJ1003, Δ*ripAF1*, CΔ*ripAF1*, and CΔ*ripAF1^2A^*. Disease severity was recorded at 15 dpi. Each time point represents the mean disease severity of 24 inoculated plants per treatment. Error bars represent the standard deviation from three independent experiments (ANOVA with Tukey’s test, \**p* < 0.05).

### RipAF1^2A^ without mADP-RT activity is unable to eliminate NbFBN1a-mediated signaling

To discover whether RipAF1 mADP-RT activity was correlated with NbFBN1a function *in vivo*, a split luciferase assay was conducted to examine the interaction affinity between RipAF1^2A^ and NbFBN1a. Compared with wild-type RipAF1, co-expression of RipAF1^2A^ and NbFBN1a remarkably reduced luciferase activity, suggesting that the interaction affinity was greatly reduced (Figure 7, A). The reduced interaction affinity was additionally verified by BiFC assays. In contrast with the strong fluorescence signal from cell membranes co-expressing nYFP-NbFBN1a and cYFP-RipAF1, an extremely weak signal was observed in the cells co-expressing nYFP-NbFBN1a and cYFP-RipAF1^2A^ (Figure 7, B). The expression of *PDF1.2* was chosen as a marker to study the effect of RipAF1 mADP-RT activity on NbFBN1a function. Overexpression of NbFBN1a showed an activation effect on *PDF1.2* promoter activity while co-expression of RipAF1 with NbFBN1a eliminated this activation (Figure 7, C and D). In contrast, co-expression of RipAF1^2A^ with NbFBN1a was unable to eliminate the activation of *PDF1.2* (Figure 7, E). The similar results were observed in NbFBN1a transgenic plants (Supplemental Figure S8, G-I). Therefore, the RipAF1^2A^ allele carrying mutations at Arg191 and Asp310 not only showed a weakened interaction with NbFBN1a, but also lost the ability to eliminate NbFBN1a-mediated signaling. This suggested that mADP-RT activity is required for RipAF1 to interfere with NbFBN1a-mediated signaling.

**Figure 7.**
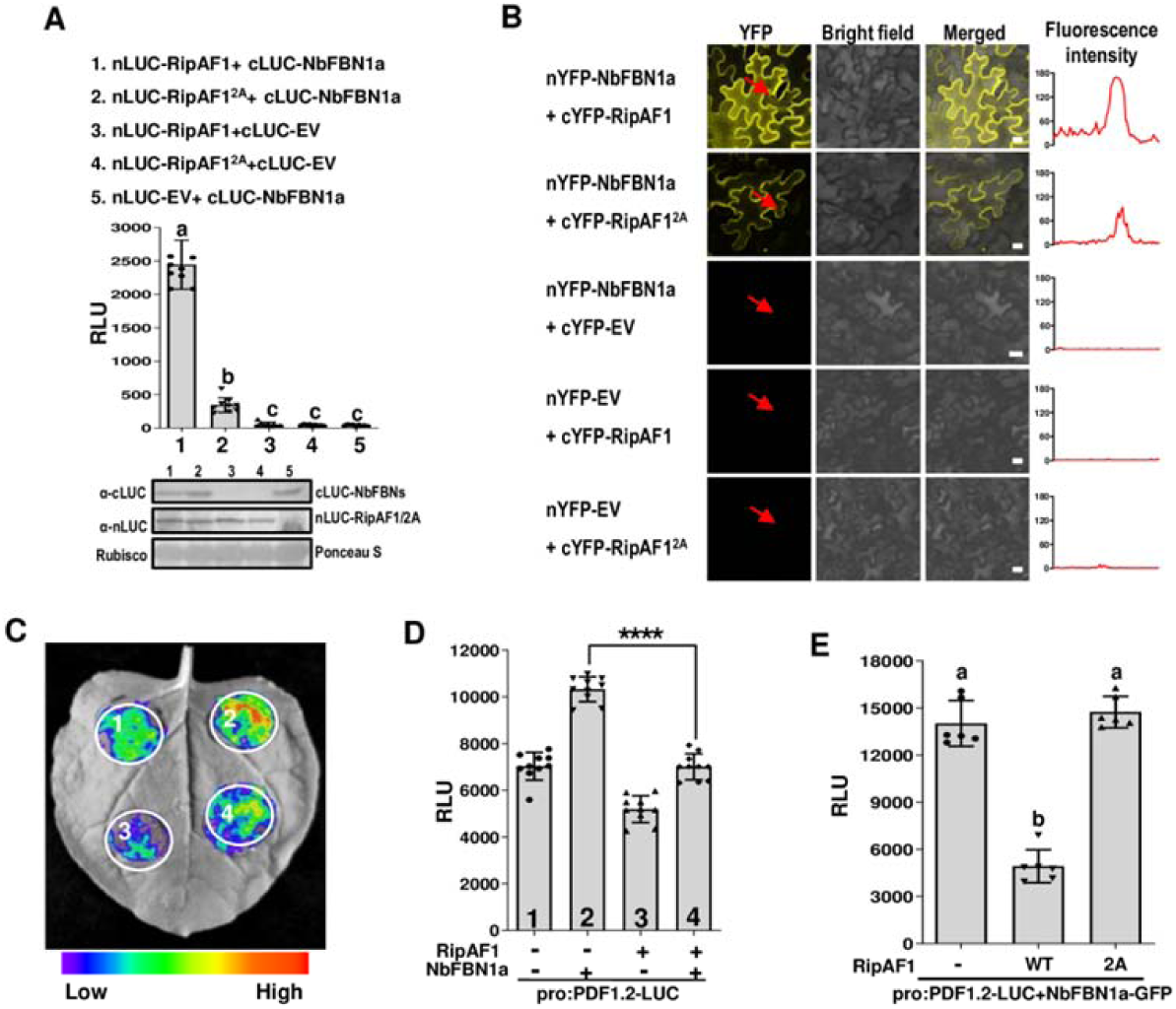
RipAF1^2A^ lost the ability to eliminate NbFBN1a-mediated signaling. **A,** Split luciferase assays to examine the reduced interaction affinity of RipAF1^2A^ with NbFBN1a. Quantification of the luciferase signal was performed with a microplate luminescence reader. Values are means ± SD (*n* = 9 biological replicates). Columns labeled with the same letter represent non-significantly different means (ANOVA with Tukey’s test, *p* < 0.01). The gels below show the expression levels of the respective proteins. **B,** BiFC assays to examine the reduced interaction affinity of RipAF1^2A^ with NbFBN1a. NbFBN1a was fused with nYFP, and RipAF1 or RipAF1^2A^ was fused with cYFP. The right panel shows the representative fluorescence intensity of the region, as indicated by arrows. The images were captured using a confocal microscope 48 h post-inoculation. Similar results were observed in three biological replicates. Scale bar, 25 μm. **C,** Wild-type RipAF1 showed the ability to eliminate NbFBN1a-induced expression of *PDF1.2. PDF1.2* promoter activity was detected by a luciferase assay. The pro:PDF1.2-Luc construct was co-transformed with RipAF1-FLAG and NbFBN1a-GFP in *N. benthamiana* leaves. Luciferase activity was measured with a CCD imaging system. **D,** Quantitative assays of the luciferase signal in C. Quantification was conducted with a microplate luminescence reader. Values are means ± SD (*n* = 10 biological replicates; Student’s *t*-test, \*\*\*\**p* < 0.0001). **E,** RipAF1^2A^ was unable to eliminate NbFBN1a-induced expression of *PDF1.2*. The analyses were performed as in D. Values are means ± SD (*n* = 6 biological replicates). Columns labeled with the same letter represent non-significantly different means (ANOVA with Tukey’s test, *p* < 0.01).

### E175/K207 are the major residues enabling NbFBN1a to be MARylated by RipAF1

To determine whether NbFBN1a was directly mono-ADP-ribosylated (MARylated) by RipAF1, GST-RipAF1 and His-MBP-tagged NbFBN1a were co-expressed in *E. coil*. By comparison with co-expressed GST tag control, all fragmentation patterns of mono-ADP-ribosylation (MARylation) were analyzed for His-MBP-NbFBN1a, including Cys, Asp, Glu, Lys, Asn, Arg, Ser, and Thr residues. E175 and K207 (E175/K207) in NbFBN1a were finally identified as the MARylation residues for RipAF1 (Figure 8, A). E175/K207 residues are highly conserved among FBN1 from *S. lycopersicum*, *S*. *tuberosum*, *C. annuum*, *A. thaliana*, and *Oryza sativa* (Supplemental Figure S9, A and B). The structure of NbFBN1a predicted by Rosetta revealed that E175/K207 residues are all located at the protein surface (Supplemental Figure S9, C), suggesting these residues can be modified by RipAF1 through protein– protein interaction. The MARylation on NbFBN1a *in vivo* was studied by expressing RipAF1-FLAG in NbFBN1a-GFP transgenic plants. The anti-panADPR immunoblot revealed a positive band after NbFBN1a-GFP fusion protein was immunoprecipitated by GFP beads (Figure 8, B). Compared with wild-type RipAF1, expression of RipAF1^2A^ in transgenic plants greatly reduced the MARylation level of NbFBN1a (Figure 8, B). Thus, RipAF1 had the ability to MARylate NbFBN1a *in vivo*. Consistent with the split luciferase and BiFC assays for the weakened interaction of RipAF^2A^ with NbFBN1a, GFP immunoprecipitation pulled less RipAF1^2A^ than wild-type RipAF1, supporting that the interaction affinity of RipAF^2A^ with NbFBN1a was reduced (Figure 8, B). To elucidate the roles of E175/K207 residues in MARylation, the two residues were mutated to alanine, thus generating NbFBN1a^E175A/K207A^ (NbFBN1a^2A^). In this case, the MARylation level of NbFBN1a^2A^ was significantly reduced relative to wild-type NbFBN1a *in vivo* (Figure 8, C). This implied that E175 and K207 in NbFBN1a are the two major residues subject to RipAF1-mediated MARylation. Even though NbFBN1a^2A^ and NbFBN1a showed similar subcellular localization to plastids and the cell membrane, the interaction affinity of NbFBN1a^2A^ with RipAF1 was reduced (Supplemental Figure S9, D and E). Notably, NbFBN1a^2A^ was impaired in activating the promoter activity of *PDF1.2* (Figure 8, D and E; Supplemental Figure S9, F). These results suggest that NbFBN1a E175/K207 are the key residues that are MARylated by RipAF1 for signaling transduction.

**Figure 8.**
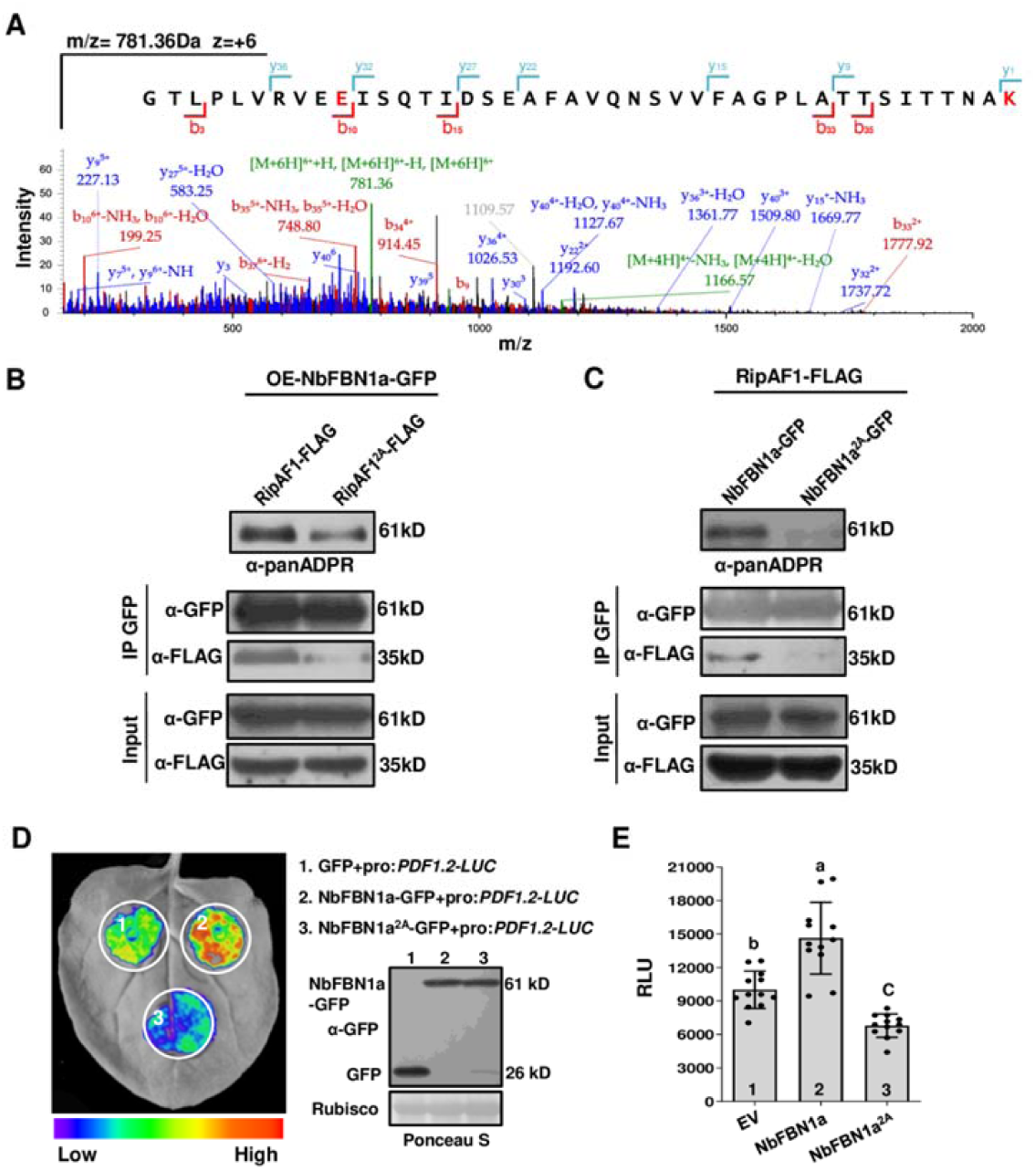
The E175/K207 residues in NbFBN1a are MARylated by RipAF1. **A,** LC-MS/MS analysis of a peptide containing NbFBN1a with MARylated E175/K207 residues. Recombinant RipAF1 and NbFBN1a proteins were co-expressed in *E. coil*. Trypsin-digested recombinant proteins were subjected to LC-MS/MS analysis. Fragment ions retaining charges in the N terminus and C terminus are denoted by “b” and “y”, respectively. **B,** NbFBN1a is mono-ADP-ribosylated (MARylated) by RipAF1 *in vivo*. RipAF1-FLAG or RipAF1^2A^-FLAG was expressed in four-week-old *NbFBN1a* transgenic plant leaves. Protein extracts were subjected to immunoprecipitation (IP) with anti-GFP agarose beads and immunoblotting (IB) with a-FLAG antibody. The ADP-ribosylation level of NbFBN1a was detected with anti-panADPR antibody. The experiment was replicated three times with similar results. **C)** The E175/K207 residues of NbFBN1a were the major residues MARylated by RipAF1. RipAF1-FLAG was co-expressed with NbFBN1a-GFP or NbFBN1a^2A^-GFP in wild-type *Nicotiana benthamiana*. In NbFBN1a^2A^, the glutamic acid at position 175 (E175) and lysine at position 207 (K207) were both substituted with alanine. Protein extracts were subjected to IP with GFP resins and IB with a-FLAG antibody. Input proteins are shown by a-FLAG IB and Ponceau S staining for protein loading (bottom). The ADP-ribosylation levels of NbFBN1a and NbFBN1a^2A^ were detected by anti-panADPR antibody. The experiment was replicated three times, with similar results. **D,** NbFBN1a^2A^ was unable to activate the expression of *PDF1.2*. The pro:PDF1.2-Luc construct was co-expressed with NbFBN1a-GFP or NbFBN1a^2A^-GFP in *N. benthamiana*. The activity of *PDF1.2* promoter was detected by a luciferase assay. The co-expression with GFP was used as the negative control. The gels at right show the expression of the respective proteins. **E,** Quantitative assays for luciferase signal. Quantification of the luciferase signal was performed with a microplate luminescence reader. Values are means ± SD (*n* = 12 biological replicates). Columns labeled with different letters represent significantly different means (ANOVA with Tukey’s test, *p* < 0.01).

## Discussion

The heterogeneous *Rso* is considered a species complex based on both phenotypic and genetic features. A prominent trait of this species complex is that it infects a wide range of hosts, with host specificity or fitness that is determined by the repertoire of Rips (Fujiwara et al., 2020; Le Roux et al., 2015; Macho et al., 2010; Xian et al., 2020). RipP1 and RipAA from GMI1000 and RipB from RS1000 induce hypersensitive response (HR) reactions and are involved in incompatibility interactions in *Nicotiana* spp. (Nakano and Mukaihara, 2019b; Poueymiro et al., 2009). The present work examined the role of RipAF1 during *Rso* infection. We found that RipAF1 plays an important role in defense induction and thereby has a negative role in bacterial virulence. Even though it belongs to *avrpPhF* family, RipAF1 does not show avirulent property on *N. benthamian* and other host plants.

SA signaling activation is required for resistance to biotrophic and hemitrophic pathogens, whereas JA signaling activation is often required for resistance to necrotrophic pathogens and herbivores (Gomi, 2020). Suppressed SA and enhanced JA signaling have been considered as disease signaling involved in bacterial wilt development (Nakano and Mukaihara, 2018). We found that RipAF1 induced the expression of the SA marker genes *ICS1*, *PR1*, and *PR2*; by contrast, the JA marker genes *PDF1.2*, *OPR3*, and *LOX* were suppressed. The changed gene expression profile suggests that JA and SA signaling were altered in host plants in response to RipAF1, resulting in a suppression effect on bacterial wilt disease signaling. Owing to the loss of this subversion effect, deletion of RipAF1 induced increased virulence and fitness of the *Rso* strains FJ1003 and GMI1000. The enhanced virulence and fitness of *ripAF1* mutants among three different plant species suggested that RipAF1 is able to induce resistance across a range of solanaceous crops.

To discover the molecular events mediated by RipAF1 in hosts, we first screened its interacting protein NbFBN1a from *N*. *benthamiana* by using the Y2H system. The interaction between RipAF1 and NbFBN1a was further verified both *in vivo* and *in vitro*. The RipAF1-GFP fusion construct in this study was observed in the cytoplasm, cell membrane, and nucleus, consistent with a report on the subcellular locations of a set of *Rso* Rips (Jeon et al., 2020). AtFBN1a was found to be located in the plastid of *Arabidopsis* (Gámez-Arjona et al., 2014). Similarly, NbFBN1a was observed in the plastid of *N. benthamiana*, but we found some FBN1a was located in the cell membrane. RipAF1 and NbFBN1a interacted at the cell membrane *in vivo*. As in *A. thaliana* (Singh et al., 2012), there are two copies of NbFBN1 in *N. benthamiana*, namely NbFBN1a and NbFBN1b, which share high similarity with homologs from various solanaceous plants. The fibrillin family is widely distributed among photosynthetic organisms (Laizet et al., 2004). In addition to its interactions with NbFBN1a/b, RipAF1 also interacts with NbFBN1 homologs from *S. lycopersicum* and *C. annuum*, implying that FBN1 homologs in the two hosts are also targeted by RipAF1.

Fibrillins represent a ubiquitous protein family that is classified into 12 phylogenetic groups (Singh and McNellis, 2011). The fibrillins in group 1 have been well documented for their biological function, especially the *Arabidopsis* fibrillins AtFBN1a and AtFBN1b (Singh and McNellis, 2011). Experimental evidence suggests that FBN1 is involved in plastoglobule formation and thylakoid maintenance (Rey et al., 2000; Simkin et al., 2007). FBN1a, FBN1b, and FBN2 interact with each other, potentially forming a network around the plastoglobule surface (Torres-Romero et al., 2022). Furthermore, FBN1 is closely related with plant hormones. In *Arabidopsis*, ABA treatment stimulated the accumulation of FBN1a protein, which in turn protected photosystem II from photoinhibition (Yang et al., 2006). RNA interference of FBN1 retarded shoot growth and reduced anthocyanin accumulation under high light combined with cold, with both effects abolished by JA treatment (Youssef et al., 2010). Knock out of *FBN1b* in *Arabidopsis* plants and knock down of *FBN1* in *Lycopersicon esculentum* plants resulted in increased susceptibility to *P. syringae* and *B. cinerea*, respectively (Leitner-Dagan et al., 2006). Through overexpression and RNA interference analysis, NbFBN1a was demonstrated to be involved in JA and SA signaling pathways. In contrast with the resistance roles of homologs in *Arabidopsis* and *L. esculentum*, NbFBN1a in *N. benthamiana* acts as a susceptibility factor for *Rso*. The *N. benthamiana* plants overexpressing NbFBN1a exhibited more severe bacterial wilting symptom, while knock-down of NbFBN1a/b enhanced resistance to bacterial wilt.

To determine how RipAF1 induces defense by targeting susceptibility factor NbFBN1a, mADP-RT activity of RipAF1 was examined, as it contains a conserved mADP-RT domain at positions 155–338. In agreement with the research on *P. syringae* pv. *tomato* HopF2 (Wang et al., 2010), the conserved Arg191 and Asp310 residues were critical for RipAF1 mADP-RT activity. By combining LC-MS/MS and site-directed mutagenesis analysis, it was found that RipAF1 directly MARylates the E175/K207 residues of NbFBN1a. Additionally, mutation of E175/K207 residues in NbFBN1a impaired its ability to modulate associated signaling. Thus, it is proposed that MARylation of NbFBN1a is the key process for RipAF1 to induce a defense reaction. As NbFBN1 has a role in bacterial wilt susceptibility, the ADP-ribosylation by RipAF1 subverts the NBFBN1a–mediated disease signaling pathways, thus promoting defense responses in host plants.

The present study was the first to report effector-mediated MARylation in the *Rso*– host interaction system. MARylation is evolutionarily conserved among all eukaryotes and prokaryotes, representing the enzymatic transfer of an ADP-ribose moiety from NAD^+^ onto target substrates with the release of nicotinamide (Feijs et al., 2013). MARylation is an important mechanism for post-translational modification of proteins that regulate a variety of cellular processes (Lüscher et al., 2018). This particular modification mode has been well documented in plant-associated *Pseudomonas syringae* effectors. In addition to the known HopF2 protein, HopU1 modifies RNA-binding proteins, including the glycine-rich RNA-binding protein GRP7, to suppress plant innate immunity in a manner dependent on its ADP-RT active residue (Fu et al., 2007). It is obvious that plant bacterial effectors with ADP-RT activity substrates not only function as host defense targets but also as susceptibility targets involved in disease development.

In contrast with HopF1 and HopF2, which are required for virulence in *P. syringae*, mutation of RipAF1 enhanced virulence of *Rso*. We suggest that this difference is explained by the different targets during bacterial infection. Introduction of HopF2 into a *hopF2* loss-of-function mutant of *P. syringae* DC3000 enhanced bacterial growth by approximately sevenfold, indicating the essential role of *hopF2* in virulence. Expression of HopF2 in *P. syringae* pv. *tabaci* induced a HR reaction in tobacco W38 plants (Wang et al., 2010). Our data showed that overexpression of *Rso* RipAF1 in *N. benthamiana* did not induce an obvious HR phenotype, while RipAF1 MARylated NbFBN1a to generate a signaling pathway antagonistic to bacterial wilt development. Accompanied with the alteration in signaling pathways, *NbFBN1-*silenced plants expressed a dwarf phenotype with decreased root length. This demonstrated that NbFBN1a/b is required for the growth of *N. benthamiana*, showing a function similar to that of AtFBN1a in *Arabidopsis* (Youssef et al., 2010).

In conclusion, we reported that the fibrillin family member NbFBN1 is involved in activating the JA signaling pathway and suppressing the SA signaling pathway in *N. benthamiana*. This is a susceptible signal transduction process that promotes bacterial wilt development. The antagonistic signaling pathways of JA and SA are disproportionate when RipAF1 is delivered into host cells. RipAF1 interacts with NbFBN1a at the cell membrane, and MARylated NbFBN1a (at the E175/K207 residues) subverts the susceptible signal pathways, thereby inducing the defense reaction (Figure 9).

**Figure 9.**
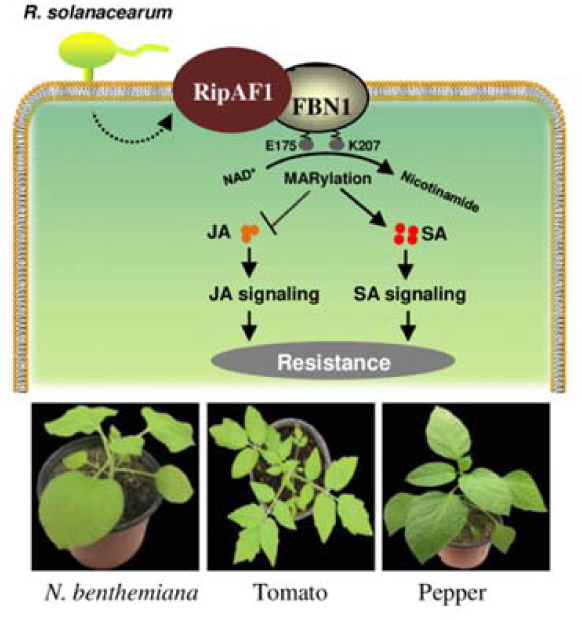
A proposed work model for the mono-ADP-ribosylation (MARylation) of FBN1 by RipAF1 that involves jasmonic acid (JA) and salicylic acid (SA) signaling. RipAF1 interacts with FBN1 at the cell membrane and MARylates FBN1 at residues E175 and K207, thus suppressing the JA signaling pathway and activating the SA signaling pathway. The MARylation process eliminates the FBN1-mediated susceptibility to promote host resistance to bacterial wilt.

## Materials and Methods

### Plant and bacterial materials

*N. benthamiana*, *S. lycopersicum* cv. ‘Hongyangli’, and *C. annuum* cv. ‘Yanshan01’ seeds were collected by the plant bacterium group at Fujian Agriculture and Forestry University (Fujian Province, China). Seeds were germinated into seedlings that were cultivated in a greenhouse under a 16/8-h light/dark photoperiod at 25°C. *Rso* FJ1003 and GMI1000 strains were used for virulence assays. The *ripAF1* mutants derived from FJ1003 and GMI1000 and the corresponding complemented strains were used to examine the contribution of RipAF1 to virulence. The complementary strains were generated by cloning *ripAF1* and its native promoter into the plasmid pBBR1MCS-5. Site-directed mutagenesis of RipAF1 to obtain RipAF1^2A^ (Arg191 and Asp310 residues were replaced with alanine) was conducted by overlap extension PCR with the primers RipAF1-R191A-F/R and RipAF1-D310A-F/R (Supplemental Table S1). The resulting constructs were verified by sequencing and introduced into the *ΔripAF1* mutant strain by electroporation.

### *Agrobacterium*-mediated transient expression

*A. tumefaciens* GV3101 cells harboring different constructs of interest were cultured in Luria-Bertani medium at 28°C and collected by centrifugation. The pellets were suspended in infiltration medium (10 mM MgCl_2_, 10 mM MES, pH 5.7, 200 μM acetosyringone) to OD_600_ = 0.3. The cell suspensions were infiltrated into *N. benthamiana* leaves using a needleless syringe. Host response, subcellular localization, and BiFC were examined 48 h post-inoculation (hpi).

### *R. solanacearum* infection assay

Bacterial virulence and plant resistance were assessed by disease severity and the percent severity index. *Rso* cells were prepared to a concentration of 1 × 10^8^ CFU/mL. Four-week-old plant stems were wounded and then inoculated with the bacterial suspension (Fan et al., 2020). Inoculated plants were maintained at 28°C for disease severity and percent severity index calculation. The percent severity index calculation was recorded on a 0-to-4 disease severity scale as follows: 0, no wilting; 1, 1–25% of leaves wilted; 2, 26–50% of leaves wilted; 3, 51–75% of leaves wilted; and 4, 76– 100% of leaves wilted. The percent severity indexes were recorded as previously described (Sun et al., 2020). All experiments were replicated three times, and each replicate contained eight plants for every strain. Bacterial growth in planta was determined as described previously (Yu et al., 2020). In brief, 100 μL of 1 × 10^6^ CFU/mL bacterial suspension was injected into four-week-old plant stems. The stem samples were collected at 1 cm above and below the injection sites 3 days post-inoculation (dpi). Bacterial growth was counted as the number of CFU per gram of stem. The experiment was conducted with three independent biological replicates.

### Quantitative RT-PCR

Total RNA was extracted by using Trizol (Invitrogen, Carlsbad, CA, USA). One microgram of total RNA was used for cDNA synthesis with HiScript QRT SuperMix with genomic DNA wipe (Vazyme, Nanjing, China). PCR was conducted on a CFX connect system (Bio-Rad, Hercules, CA, USA) with ChamQ SYBR qPCR Master Mix (Genstar, Beijing, China). *EF1α* was used as an internal control. Primers for target genes are listed in Supplemental Table S1. The PCR thermal cycling conditions consisted of denaturation at 95°C for 5 min, followed by 40 cycles of DNA denaturation at 95°C for 10 s and annealing at 58°C for 20 s. All the experiments included three biological replicates, each with three technical replicates.

### Yeast two-hybrid assays

Yeast two-hybrid assays were conducted to screen for RipAF1-interacting proteins within the cDNA library of *N. benthamiana* according to a standard AcLi-mediated transformation kit (Matchmaker; Clontech, Mountain View, CA, USA). To repeat the interaction, the coding sequences of *RipAF1* and *NbFBN1a* were individually cloned into both pGADT7-T and pGBKT7 vectors. AD-RipAF1/BD-NbFBN1a or AD-NbFBN1a/BD-RipAF1 was then transformed into yeast strain AH109 and screened on synthetic drop-out media lacking leucine and tryptophan (SD/-Leu-Trp). Single colonies were cultured, serially diluted, and grown on SD/-Leu-Trp and SD/-Leu-Trp-His media to examine the possible interaction. Then, 1 mM 3-amino-1,2,4-triazole (3-AT) was supplied to SD/-Leu-Trp-His to inhibit the autoactivation of AD-NbFBN1a when AD-NbFBN1a/BD-RipAF1 was co-transformed into yeast. Additionally, pGADT7-T and pGBKT7-53 were used as positive controls, while pGADT7-T and pGBKT7-lam served as negative controls.

### MBP pull-down assay

*NbFBN1a* was cloned into pMAL-C4X to express MBP-NbFBN1a fusion protein. *RipAF1* was cloned into pGEX-4T to express GST-RipAF1 fusion protein. MBP-NbFBN1a and GST-RipAF1 were separately expressed in *E. coli* BL21(DE3) by induction with 1.0 mM isopropyl-β-D-thiogalactopyranoside (IPTG). After the proteins were purified by glutathione resin and amylose affinity chromatography, MBP pull-down assays were performed (Wu et al., 2019).

### Luciferase assay and detection

The split-luciferase complementation assay was performed to verify the interaction between RipAF1 and host FBN1 *in vivo* (Chen et al., 2008). *FBN1* genes from *N. benthamiana*, *S. lycopersicum*, and *C. annuum* were individually fused with cLUC at the C terminus. Each cLUC-FBN1 construct was co-expressed with nLUC-RipAF1 in 4-week-old *N. benthamiana* leaves. To examine the role of conserved residues involved in the putative interaction, RipAF1^2A^ was fused with nLUC to generate nLUC-RipAF1^2A^. NbFBN1a^2A^ (E175/K207 residues were replaced by alanine) was fused with cLUC by overlap extension PCR with the primers NbFBN1a-E175A-F/R and NbFBN1a-K207A-F/R (Supplemental Table S1) to generate cLUC-NbFBN1a^2A^. For *PDF1.2* promoter activity analysis, a 1493-bp promoter region in the pDONR207 vector was directly introduced into the destination vectors pGWB435 via Gateway cloning to generate pro:*PDF1.2*-LUC. *PDF1.2* promoter activity was detected after transient expression in *N. benthamiana.* For each detection assay, the leaves were collected 48 hpi, rubbed with 0.5 mM luciferin, and kept in the dark for 1 min to quench the fluorescence. A cooled CCD imaging apparatus (Roper Scientific, Trenton, NJ, USA) was used to capture luciferase luminescence images. The luciferase signal strength was quantified to monitor interaction affinity and promoter activity (Yu et al., 2020). Briefly, the leaf discs were incubated with 100 μL of water containing 0.5 mM luciferin in a 96-well plate, and the luminescence was recorded with a microplate luminescence reader (Varioskan Flash, Thermo Scientific, Waltham, MA, USA).

### Subcellular localization analysis

Subcellular localization of every fluorescence fusion construct was examined with a confocal laser microscope (Leica Model TCS SP8; Leica, Wetzlar, Germany) at 48 h post-transient expression in *N. benthamiana.* RipAF1-GFP and NbFBN1a/b-GFP were observed at a 488 nm excitation wavelength with a 470–550 nm bandpass emission filter. The subcellular localization of NbFBN1a/b-GFP was additionally examined in the protoplast of *N. benthamiana* (Yoo et al., 2007). For co-localization analysis, RipAF1 was cloned into pGD-3G-mCherry (Sun et al., 2018), generating a RipAF1-RFP fusion construct. RipAF1-RFP was co-expressed with GFP-NbFBN1a/b and observed at a 514 nm excitation wavelength with a 530–560 nm bandpass emission filter. Both pSCYNE and pSCYCE plasmids were used for the BiFC assays (Waadt et al., 2008). RipAF1 was cloned into pSCYCE at the C terminus, generating the cYFP-RipAF1 construct. NbFBN1a/b were cloned into pSCYNE at the N terminus, generating NbFBN1a/b-nYFP constructs. cYFP-RipAF1^2A^ and NbFBN1a^2A^-nYFP were additionally constructed for BiFC assays. Yellow fluorescence was observed at a 514 nm excitation wavelength with a 500–550 nm band-pass emission filter. The membrane-bound marker PIP2A-mcherry was applied in the BiFC assay to verify membrane localization (Nelson et al., 2007). Every subcellular localization analysis was replicated three times.

### Generation of *NbFBN1a-GFP* transgenic plants

To generate *NbFBN1a-GFP* overexpressing plants, the *NbFBN1a* coding sequence was cloned into the pCAMBIA1300s-GFP binary vector driven by a CaMV 35S promoter. The construct was verified by sequencing and then used for plant transformation via the leaf-disc method after introduction into *A. tumefaciens* strain HB101. The *NbFBN1a-GFP* overexpressing plants were selected with 25 μg mL^-1^ hygromycin. The primer pair NbFBN1a-GFP-F and NbFBN1a-GFP-R was used for PCR amplification to verify transgene integration in T0 and T1 offspring. T2 transgenic homozygous lines were used for gene expression and disease resistance analysis. The expression of NbFBN1a-GFP was verified by western blot with α-GFP antibody.

### Phytohormone measurement

Phytohormones were quantified by Nanjing Convinced-Test Technology Company (Nanjing, China) using HPLC-MS/MS. Leaves (150 mg in fresh weight) were homogenized in liquid nitrogen and extracted twice at 4℃ for 1 h using an extraction solvent of isopropanol-water-hydrochloric acid (2:1:0.002, *V*:*V*:*V*). The extracted samples were spiked with H_2_JA (Tokyo Chemical Industry, Tokyo, Japan) and d4-SA (OlchemIm, Olomouc, Czech Republic) as internal standards. Phytohormones were then measured with an Agilent 1290 HPLC device (Agilent Technologies, Santa Clara, CA, USA) with an AB SCIEX QTRAP 6500 MS/MS system (AB SCIEX, Framingham, MA, USA). Experiments were performed with three independent biological replicates.

### Virus-induced gene silencing

A 300-bp partial sequence of *NbFBN1a* was inserted into the pTRV2 vector to knock down *NbFBN1a/b* in *N. benthamiana*; the pTRV2:*NbPDS* construct was used as the control to evaluate silencing efficiency. The pTRV2:*gfp* construct carrying a 358-bp fragment of the *gfp* gene was used as the negative control (Sun et al., 2020). A mixture of GV3101 cultures (1:1, *v*/*v*) containing pTRV1 and pTRV2 constructs was co-infiltrated into 20-day-old *N. benthamiana* leaves. When *NbPDS*-silenced plants showed a photo-bleached phenotype, qRT-PCR was performed to detect the *NbFBN1a/b* transcript levels. *NbFBN1a/b*-silenced plants were evaluated for signaling marker gene expression, phytohormone contents, and bacterial wilt resistance.

### Co-immunoprecipitation

Co-immunoprecipitation (Co-IP) was performed on homozygous *NbFBN1a-GFP* transgenic plants according to a previous method (Luo et al., 2017). At 48 hpi transient expression of RipAF1-FLAG in transgenic plants, 1.5-g samples of inoculated leaves were ground in liquid nitrogen and extracted in extraction/washing buffer containing 50 mM Tris-HCl (pH 7.5), 150 mM NaCl, 0.1% Triton-100, 0.2% Nonidet P-40, 1 mM DTT, 1×complete protease inhibitor (Roche, Shanghai, China), and 1×phosphatase tablet (Roche). The homogenate was centrifuged at 15,000 × *g* for 20 min. Anti-GFP antiserum-conjugated agarose beads (Sigma, Shanghai, China) were added to the supernatant. After incubation at 4°C with end-over-end shaking for 1.5 h, the beads were spun down at 1,500 × *g* for 2 min and washed with washing buffer at least five times. The bound proteins were eluted by 1.5× Laemmli loading buffer, resolved by 10% SDS-PAGE, and subjected to western blotting analysis.

### Western blotting analysis

Western blotting was performed to detect protein expression in *N. benthamiana* according to a previously described method (Luo et al., 2017). Briefly, leaf samples were harvested at 48 hpi. The plant leaves were ground in liquid nitrogen and extracted in extraction/washing buffer (Roche). The extraction proteins were eluted with 1.5×Laemmli loading buffer, resolved by 10% (*w*/*v*) SDS-PAGE, and subjected to western blotting analysis using anti-FLAG (1:10000) or anti-GFP polyclonal antibody (1:5000).

### LC-MS/MS analysis of ADP-ribosylation residues

To identify the residues of NbFBN1a mono-ADP-ribosylated by RipAF1, GST-RipAF1 and MBP-NbFBN1a were co-expressed in *E. coli* strain BL21. MBP-NbFBN1a was purified by amylose affinity chromatography. Ten micrograms of MBP-NbFBN1a protein were incubated with 3×Laemmli loading buffer at 95°C for 5 min. Denatured proteins were separated by 8% SDS-PAGE and subjected to trypsin digestion. Phosphopeptides were analyzed by the Q Exactive LC-MS/MS system (Thermo Scientific, Beijing Qinglian Biotech Co., Ltd, Beijing, China). Acquired mass spectra were analyzed using MaxQuant software, and the phosphorylation sites were manually verified. The auto-mono-ADP-ribosylation residues in RipAF1 were identified by applying 10 μg of purified GST-RipAF1 proteins from *E. coli* strain BL21 harboring a GST-RipAF1 construct.

### Detection of ADP-ribosylation activity

An anti-panADPR (catalog no. MABE1016, EMD Millipore, Burlington, MA, USA) was used to detect ADP-ribosylated proteins. GST-RipAF1 or GST-RipAF1^2A^ was expressed in *E*. *coli* strain BL21(DE3) and purified with a glutathione resin. To detect the ADP-ribosylation of RipAF1 *in vivo*, RipAF1-GFP and RipAF1^2A^-GFP were transiently expressed in *N. benthamiana* plants and purified by using GFP-Trap_A (ChromoTek, Planegg, Germany). The bound proteins were eluted with 3×Laemmli loading buffer, resolved by 10% SDS-PAGE, and then subjected to ADP-ribosylation analysis using a 1:5000 (*v*/*v*) dilution of anti-panADPR.

To detect ADP-ribosylated NbFBN1a, RipAF1-FLAG was first transiently expressed in *NbFBN1a-GFP* transgenic plants. The expression of RipAF1^2A^-FLAG was used as a negative control. NbFBN1a-GFP proteins were pulled down by Co-IP using GFP-Trap_A (ChromoTek) for ADP-ribosylation analysis. To verify the involvement of E175/K207 residues in ADP-ribosylation, RipAF1-FLAG and NbFBN1a^2A^-GFP were transiently co-expressed in *N. benthamiana* plants. The co-expression of RipAF1-FLAG and NbFBN1a-GFP was used as the positive control. The ribosylated NbFBN1a^2A^-GFP and NbFBN1a-GFP were detected by the anti-pan-ADP-ribose binding reagent after they were pulled down with Co-IP by GFP-Trap_A (ChromoTek).

### Sequence analysis

The distribution of RipAF1 in diverse phylotypic *Rso* strains was collected from the Ralsto T3E database (https://iant.toulouse.inra.fr/bacteria/annotation/site/prj/T3Ev3/). The RipAF1 and fibrillin sequences were downloaded from the UniProt database (http://uniprot.org). Sequence alignments were generated using Clustal X with default settings and analyzed by Espript3.0 (http://espript.ibcp.fr/ESPript/cgi-bin/ESPript.cgi). The phylogenetic tree was generated from amino acid sequences using MEGA 7.0 (Kumar et al., 2016). The bootstrap consensus tree inferred from 1000 replicates is taken to represent the evolutionary history of the taxa analyzed. The consensus amino acid positions of *Rso* RipAF1 and host FBN1 were generated using WebLogo (http://weblogo.threeplusone.com/create.cgi). Three-dimensional structures were predicted by Rosetta and RaptorX (http://raptorx.uchicago.edu).

### Quantification and statistical analysis

All statistical analyses were conducted by using the GraphPad Prism software (GraphPad Software, San Diego, CA, USA) with an unpaired, two-tailed Student’s *t*-tests or one-way ANOVA at the 95% level with Tukey’s multiple comparison test. The values are presented as means ± standard deviation (SD).

## Supplemental data

Supplemental Figure S1. RipAF1 with a mono-ADP-ribosyltransferase domain was conserved in *Ralstonia solanacearum*.

Supplemental Figure S2. RipAF1 mutant of GMI1000 *Ralstonia solanacearum* showed enhanced virulence to *Solanum lycopersicum* and *Capsicum annuum*.

Supplemental Figure S3. RipAF1 interacts with NbFBN1a/b at the cell membrane.

Supplemental Figure S4. RipAF1 interacted with FBN1 from *Solanum lycopersicum* and *Capsicum annuum*.

Supplemental Figure S5. Transient overexpression of NbFBN1a activated jasmonic acid (JA) signaling and suppressed salicylic acid (SA) signaling in *Nicotiana benthamiana*.

Supplemental Figure S6. Detection of *NbFBN1a* expression in transgenic *Nicotiana benthamiana* plants.

Supplemental Figure S7. *PDF1.2* promoter activity was reduced in *NbFBN1a-*silenced *Nicotiana benthamiana*.

Supplemental Figure S8. The conservation of R191/D310 residues among RipAF1 homologs.

Supplemental Figure S9. E175/K207 residues were necessary for NbFBN1a function.

Supplemental Table S1. List of primers used in this work.

## Supplemental data

**Figure S1. RipAF1 with a mono-ADP-ribosyltransferase domain was conserved in *Ralstonia solanacearum*.**

**A,** Distribution of RipAF1 in representative *R. solanacearum* species from four phylotypes. Green indicates one copy in the species, while red indicates the effector is missing. **B,** Phylogenetic analysis of RipAF1 from FJ1003 and its homologs from other *R. solanacearum* species. The phylogeny was inferred using the maximum likelihood method implemented in MEGA 7.0. The bootstrap consensus tree inferred from 1000 replicates represents the evolutionary history of the taxa analyzed. **C,** Schematic diagram of the RipAF1 structure possessing a mono-ADP-ribosyltransferase (mADP-RT) domain. The mADP-RT domain from 155 to 338 residues is shown in red. **D,** Amino acids sequence alignment of the mADP-RT domain in RipAF1 from FJ1003 and GMI1000, respectively, and HopF1 and HopF2 from *Pseudomonas syringae*. The sequences were aligned with Clustal X and analyzed using Espript3.0. The conserved amino acids are indicated in red. **E,** Transient expression of RipAF1 induced chlorosis in *N*. *benthamian*a. A. *tumefaciens* strains carrying pJL12 empty vector and pJL12:RipAF1 were injected into *N*. *benthamiana* leaves. Images were captured 48 h after inoculation. The infiltration areas are indicated by the dotted-line circle. The expression of RipAF1 was verified by a western blot using α-FLAG antibody.

**Figure S2. RipAF1 mutant of GMI1000 *Ralstonia solanacearum* showed enhanced virulence to *Solanum lycopersicum* and *Capsicum annuum*.**

**A,** Disease symptoms on *S. lycopersicum* stem tissue near inoculation sites inoculated with wild-type GMI1000, mutant Δ*ripAF1*, and the complemented strain CΔ*ripAF1.* Images were captured 7 d post-inoculation (dpi). All experiments were replicated three times with similar results, and representative results are shown. **B,** Progression of bacterial wilt on *S. lycopersicum* plants inoculated with wild-type GMI1000, Δ*ripAF1*, and CΔ*ripAF1.* The disease index was recorded at 7 dpi. Each time point represents the mean disease severity of 24 inoculated plants per treatment. Error bars represent the standard deviation across three independent experiments (ANOVA, \**p* < 0.05). **C,** Growth of the wild-type GMI1000, Δ*ripAF1*, and CΔ*ripAF1* strains in *S. lycopersicum* plants. The four-week-old *S. lycopersicum* plants were injected with 100 μL of 10^6^ CFU/mL bacterial cell suspension. Plants were subjected to growth curve analysis at 3 dpi. Values are means ± SD (*n* = 6 biological replicates; Student’s *t*-test, \*\*\**p* < 0.001, \*\*\*\**p* < 0.0001). **D,** Disease symptoms on *C. annuum* stem tissue near inoculation sites inoculated with wild-type GMI1000, mutant Δ*ripAF1*, and CΔ*ripAF1*. The disease index was recorded at 10 dpi. The analyses were performed as in A. **E,** Progression of bacterial wilt on *C. annuum* plants. The analyses were performed as in B. **F,** Growth of the wild-type GMI1000, Δ*ripAF1*, and CΔ*ripAF1* strains in *C. annuum* plants. The analyses were performed as in C (Student’s *t*-test, \*\*\**p* < 0.001, \*\*\*\**p* < 0.0001).

**Figure S3. RipAF1 interacts with NbFBN1a/b at the cell membrane.**

**A,** Growth of yeast co-transformed with BD-RipAF1 and AD-NbFBN1a on synthetic dextrose (SD) plates. The transformants were screened on SD media lacking leucine and tryptophan (SD/-Leu-Trp). The single yeast colonies were serially diluted and grown on SD/-Leu-Trp and SD/-Leu-Trp-His (SD media lacking leucine, tryptophan, and histidine) to examine the putative interaction. One mM 3-amino-1,2,4-triazole (3-AT) was supplied to inhibit the autoactivation of AD-NbFBN1a. Yeast co-transformed with pGADT7-T and pGBKT7-53 served as the positive control (+), and yeast co-transformed with pGADT7-T and pGBKT7-lam served as the negative control (-). EV, empty vector. **B,** Amino acid sequence alignment of NbFBN1a and NbFBN1b. The sequences were aligned with Clustal X. **C,** Subcellular localization of RipAF1-GFP and NbFBN1a/b-GFP. RipAF1-GFP and NbFBN1a/b-GFP fusions were transiently expressed in *N*. *benthamiana* leaves following *Agrobacterium*-mediated transformation. The images were captured using a confocal microscope. Scale bar, 25 μm. The arrows indicate the small amount of NbFBN1a and NbFBN1b localized to the cell membrane. **D,** NbFNB1a/b is a plastid-localized protein in the protoplast of *Nicotiana benthamiana*. Plasmids containing *35S:GFP* and *35S:NbFBN1a/b-GFP* were transformed into *N. benthamiana* protoplasts. Confocal micrographs were obtained 16 h post-transformation. Scale bar, 10 μm. **E,** NbFNB1a/b and RipAF1 were co-localized to the cell membrane in *N. benthamiana*. NbFNB1a/b-GFP and RipAF1-mcherry were co-expressed in *N. benthamiana* and visualized by confocal microscopy. GFP co-expressed with RipAF1-mcherry and NbFBN1a/b-GFP co-expressed with mcherry-EV were used as the negative controls. Scale bar, 10 μm. **F,** NbFBN1a/b interacted with RipAF1 in cell membranes of *N. benthamiana* leaves. NbFBN1a/b was fused with nYFP, and RipAF1 was fused with cYFP. The PIP2A-mcherry construct was used as a plasma membrane marker. Images were captured using a confocal microscope at 48 hpi. Scale bar, 10 μm.

**Figure S4. RipAF1 interacted with FBN1 from *Solanum lycopersicum* and *Capsicum annuum*.**

**A,** The phylogeny based on FBN1 sequences from *Nicotiana benthamiana* (*Nb*)*, Solanum lycopersicum* (*Sl*), *Capsicum annuum* (*Ca*), and *Arabidopsis thaliana* (*At*). The phylogenetic tree was generated using MEGA 7.0. The star indicates NbFBN1a from *N*. *benthamiana*. **B,** Split luciferase assay to assess the interaction of RipAF1 with SlFBN1 and CaFBN1. The protein interaction strength was quantified according to the luminescence signals. Values are means ± SD (*n* = 6 biological replicates; Student’s *t*-test, \*\*\*\**p* < 0.0001).

**Figure S5. Transient overexpression of NbFBN1a activated jasmonic acid (JA) signaling and suppressed salicylic acid (SA) signaling in *Nicotiana benthamiana.***

A, NbFBN1a activated the expression of JA marker genes. Total RNA was isolated from leaves 48 h after agroinfiltration. Expression levels were determined by qRT-PCR analysis and normalized to that of the GFP control. Values are means ± SD (*n* = 3 biological replicates; Student’s *t*-test, \**p* < 0.05, \*\**p* < 0.01). B, NbFBN1a suppressed the expression of SA marker genes, based on the same procedure as in A. Values are means ± SD (*n* = 3 biological replicates; Student’s *t*-test, \*\**p* < 0.01). C, Luciferase assay for *PDF1.2* promoter activity induced by NbFBN1a. The pro:PDF1.2-Luc and NbFBN1a-GFP constructs were co-expressed in *N. benthamiana* leaves. Luciferase activity was measured with a CCD imaging system. The co-expression of pro:PDF1.2-Luc and GFP served as the negative control. The gels at the right show the expression of respective proteins. D, Quantitative assay for luciferase signal with a microplate luminescence reader. Values are means ± SD (*n* = 6 biological replicates; Student’s *t*-test, \**p* < 0.05).

**Figure S6. Detection of *NbFBN1a* expression in transgenic *Nicotiana benthamiana* plants.**

**A,** Quantitative reverse transcription PCR analysis of the *NbFBN1a* transcript level in transgenic plants. Total RNA was isolated from four-week-old transgenic plants. The transcript level in wild-type *N*. *benthamiana* plants was used as a control to monitor expression changes. OE-25 and OE-29 are independent transgenic lines. Values are means ± SD (*n* = 3 biological replicates; Student’s *t*-test, \*\**p* < 0.01). **B,** Western blot analysis of overexpression of NbFBN1a. The NbFBN1a-GFP transgenic lines OE-25 and OE-29 were assayed by anti-GFP immunoblotting. Wild-type *N*. *benthamiana* served as the negative control. **C,** Examination of plastid-localized NbFBN1a-GFP in transgenic plant leaves. The images were captured using a confocal microscope. Scale bar, 25 μm.

**Figure S7. *PDF1.2* promoter activity was reduced in *NbFBN1a-*silenced *Nicotiana benthamiana*.**

**A,** Quantitative reverse transcription PCR analysis of the transcript level of *NbFBN1a* in *NbFBN1a*-silenced plants. RNA was isolated from the new upper leaves when photobleaching was observed in phytoene desaturase (*PDS*)-silenced positive control plants. The transcript level in plants transformed with pTRV:*gfp* was used as a control to monitor expression changes. Error bars represent the standard deviation from three replicates. Values are means ± SD (*n* = 3 biological replicates; Student’s *t*-test, \*\**p* < 0.01). **B,** The growth of plants transformed with pTRV:*NbFBN1.* The plants were collected at 3 weeks post-agroinfiltration. **C,** Luciferase assay for *PDF1.2* promoter activity in *NbFBN1a-*silenced plants. The pro:PDF1.2-Luc construct was transformed into the leaves of plants transformed with pTRV:*gfp* or pTRV:*NbFBN1*. Quantification of the luciferase signal was performed with a microplate luminescence reader. Values are means ± SD (*n* = 6 biological replicates; Student’s *t*-test, \*\**p* < 0.01).

**Figure S8. The conservation of R191/D310 residues among RipAF1 homologs.**

**A,** WebLogo analysis of the conservation of the R191 residue in RipAF1 from *Ralstonia solanacearum.* Sequences were downloaded from Swiss-Prot and aligned with Clustal X. Aligned sequences were subjected to online WebLogo analysis. The conserved arginine acid (R190) is highlighted in red. **B,** Multiple sequence alignment of the 22-amino acid domain including the R191 residue in RipAF1 homologs. Sequences of RipAF1 were downloaded from Swiss-Prot. Different pathovars of *Pseudomonas syringae* and other species from the genus *Pseudomonas* were selected for multiple sequence alignment by Clustal X. The conserved arginine (R191) is highlighted in red. **C,** WebLogo analysis of the conservation of D310 in RipAF1 from *R. solanacearum.* The analyses were performed as in A. **D,** Multiple sequence alignment of the 33-amino acid domain including the D310 residue in RipAF1 homologs. The conserved aspartic acid (D310) is highlighted in red. Other invariant residues are shown in blue. The analyses were performed as in B. **E,** The locations of R191 and D310 residues in the predicted 3D structure of RipAF1, as predicted using Rosetta. **F,** The subcellular localization of RipAF1^2A^. RipAF1^2A^-GFP was transiently expressed in *Nicotiana benthamiana* leaves following *Agrobacterium*-mediated transformation. The fluorescence was visualized by confocal microscopy. GFP and RipAF1-GFP were used as controls. Scale bar, 25 μm. **G,** RipAF1^2A^ exhibited a reduced ability to inhibit expression of *PDF1.2* on *FBN1a* transgenic plants. The activity of the *PDF1.2* promoter was detected by luciferase assay. The pro:PDF1.2-Luc construct was co-transformed with RipAF1-FLAG and RipAF1^2A^-FLAG. Luciferase activity was measured with a CCD imaging system. **H,** Quantitative assays for luciferase signal in G. Quantification was conducted with a microplate luminescence reader. Values are means ± SD (*n* = 8 biological replicates). Columns labeled with different letters represent significantly different means (ANOVA with Tukey’s test, *p* < 0.01). **I,** Western blot analysis of respective proteins in G.

**Figure S9. E175/K207 residues were necessary for NbFBN1a function.**

**A,** WebLogo analysis of the conservation of E175/K207 in FBN1 from different plant species. FBN1 sequences were downloaded from Swiss-Prot and aligned with Clustal X. Aligned sequences were subjected to online WebLogo analysis. The conserved glutamic acid (E175) and lysine (K207) residues are highlighted in red. **B,** Sequence alignments of the conserved E175/K207 residues in NbFBN1 with homologs from *Solanum lycopersicum* (*Sl*), *Solanum tuberosum* (*St*), *Capsicum annuum* (*Ca*), *Arabidopsis thaliana* (*At*), and *Oryza sativa* (*Os*). FBN1 sequences were downloaded from Swiss-Prot and aligned with Clustal X. The conserved tyrosine residue is highlighted in red. **C,** The locations of E175 and K207 residues in the 3D structure of NbFBN1a (from 92 to 321 amino acids), as predicted by Rosetta, with the locations of E175 and K207 indicated by arrows. **D,** BiFC assays of the weakened interaction of NbFBN1a^2A^ with RipAF1. NbFBN1a^2A^ was fused with nYFP, and RipAF1 was fused with cYFP. The right panel shows the fluorescence intensity, with the region of interest indicated by the arrow. The images were captured using a confocal microscope at 48 hpi. Similar results were observed in three biological replicates. Scale bar, 25 μm. **E,** Subcellular localization of NbFBN1a^2A^. NbFBN1a^2A^-GFP was expressed in *Nicotiana benthamiana* by *Agrobacterium*-mediated transient expression. NbFBN1a-GFP and GFP were used as controls. Images were obtained by confocal microscopy at 48 h post-infiltration. Scale bar, 25 μm. **F,** Quantitative assays of *PDF1.2* promoter activity co-expressed with NbFBN1a^2A^. Quantification of the luciferase signal was performed with a microplate luminescence reader. Columns labeled with the same letter indicate means are not significantly different, and the values are means ± SD (*n* = 6 biological replicates; ANOVA with Tukey’s test, *p* < 0.01).

